# Longitudinal serum proteomics analyses reveal early diagnostic and discriminatory markers for porcine influenza and coronavirus infections

**DOI:** 10.64898/2026.04.21.719833

**Authors:** Cecile Frampas, Basudev Paudyal, Juanjuan Guo, Kristien van Reeth, Anthony D. Whetton, Yashwanth Subbannayya, Elma Tchilian, Sneha M. Pinto

## Abstract

Respiratory virus infections affect both humans and livestock, causing considerable mortality and morbidity. While respiratory pathogens such as swine influenza A virus (pH1N1) and porcine respiratory coronavirus (PRCV) often present with overlapping clinical symptoms, their pathological trajectories and outcomes differ. Given the propensity for pathogen spillover and the use of pigs as a physiologically relevant large-animal translational model, we aimed to characterise host serum protein signatures that detect and differentiate pH1N1 from PRCV, enabling improved disease monitoring and control. Using high-resolution mass spectrometry-based proteomics, we identified 162 serum proteins that were significantly dysregulated across 3 infection timepoints (1, 5, and 12 days post-infection (DPI)), with signatures correlating with viral shedding and lung pathology as early as 1 DPI. Notably, multiplexed targeted analysis of a subset of proteins in an independent cohort from a different breed and geographic location demonstrated detection, femtomole-level targeted quantitation, and validation of SRGN as a diagnostic marker for pH1N1 and PRCV (AUC=0.85). Further, SOD1 was validated as an early marker for PRCV, increasing as early as 1 DPI (AUC= 0.9). Finally, a multi-peptide signature composed of SRGN, SOD1, and RAN demonstrated reasonable predictive power for pH1N1 (AUC=0.75) and PRCV (AUC=0.65) at 1 DPI. Our data validate the proteomic screening, provide insights into the role of early protein markers in distinguishing respiratory viral infections, and pave the way for the development of point-of-care diagnostics and targeted prevention strategies, enhancing preparedness against emerging zoonotic threats.

## Background

Respiratory viruses such as influenza and coronaviruses (CoV) significantly impact the human-animal-environment interface, affecting global health security and food systems (1, 2). These viruses often originate from animal reservoirs and cross species barriers, posing a risk of emerging infections, as evidenced by the influenza and COVID-19 pandemics (3–5). Understanding host responses to respiratory virus infection and identifying early predictive markers are, therefore, essential for informing control strategies and developing interventions to prevent zoonotic spillover.

Influenza and respiratory CoV infections share features such as epithelial damage and immune responses, but differ in tissue tropism, receptor usage, and symptoms (6, 7). These infections have been studied using various animal models, including mice, ferrets (8, 9), hamsters (10), and non-human primates (11). However, these are not natural hosts, and the translatability of findings to human infection is limited. Among livestock species, pigs are not only economically important as a food source but also serve as a physiologically relevant large-animal model for human-based studies (12). They are natural hosts for influenza viruses and porcine respiratory coronavirus (PRCV), which recapitulate important features of human influenza and CoV infections (13–17). Influenza A and porcine respiratory coronavirus are enzootic in pigs, with pH1N1 being zoonotic and pandemic-prone, while PRCV has no pandemic potential (15, 18). Although both pH1N1 and PRCV infections often present with overlapping aetiologies, a recent study demonstrated fundamental differences in coronavirus and influenza virus-host interactions in their large natural host, the pig (19).

We hypothesised that influenza and PRCV would elicit both shared and virus-specific temporal serum proteome responses associated with disease severity, and that these host-response signatures could support discrimination between viral aetiologies in a natural large-animal model with high relevance to human infection. Current approaches are pathogen-centric, relying primarily on virus isolation, antigen detection, and serology (20), and require targeted respiratory tract sampling and therefore can be invasive, labour-intensive, and logistically challenging to implement on a large scale or longitudinally in field conditions (21). Moreover, accurate diagnosis often relies on post-mortem examination, limiting applicability for longitudinal or large-scale field monitoring (22). Because viral loads can decline below detection thresholds even while lung pathology and systemic host responses persist, there is a critical need for host-response protein signatures that enable early, reliable differentiation of viral aetiologies. Early, farm adaptable diagnostics that can both detect infection early and discriminate between respiratory viruses are essential to support early intervention strategies and thereby minimise impact or prevent their spread (23–25).

Blood protein markers are routinely used for clinical diagnosis owing to their ease of collection and their ability to provide a snapshot of the host/immunological response to infections. With recent advances in depth, throughput, and reproducibility, mass spectrometry-based approaches for blood proteomics hold promise for identifying early, discriminating host markers that aid diagnosis, prognosis, and theranostics (26–28). Understanding changes in the proteome associated with specific virus-induced disease trajectories will inform management strategies and may help mitigate the emergence of future viral infections (29, 30). While temporal changes in the plasma/serum proteome have been well documented in human COVID-19 hospitalised cases, longitudinal sampling that enables detection of early responses and progression, and allows assessment of similar or divergent responses to other respiratory viruses, remains underexplored. Furthermore, a similar scale of studies in economically important and physiologically relevant animals is lacking (30).

In this study, we examined temporal changes in the serum proteome of a porcine natural host model following infection with pH1N1 and PRCV, using a high-throughput liquid chromatography mass spectrometry (LC-MS/MS)–based proteomics platform. We further demonstrate that discovery studies, quantitation, and targeted validation of select candidates in an independent cohort can be performed on a single instrument, thereby uncovering markers that correlate with virulence and pathogenesis. Together, our analyses provide robust insights into alterations in the serum proteome associated with disease severity and into shared and divergent mechanisms of respiratory viral pathogenesis.

## Materials and methods

### Cohort design and sample characteristics

Serum samples for the discovery phase were collected from an infectious study at the Pirbright Institute, UK, following approval by the ethics review process under the Animals (Scientific Procedures) Act 1986, with project licences PP7764821 and PP2064443. The study received a favourable retrospective governance review from the University of Surrey NASPA, a sub- committee of the Animal Welfare and Ethical Review Board (NASPA-2526-08(L)). Thirty female pigs aged 6–8 weeks were obtained from a commercial high-health status herd. The animals obtained were outbred and genetically composed of one-quarter Large White, one-quarter Landrace, and one-half Hampshire breeds. Prior to the study, pigs were screened using ELISA assays to confirm the absence of serum antibodies against the PRCV spike protein and the haemagglutinin (HA) of H1N1pdm09, H1N2, H3N2, and avian-like H1N1. Animals were then randomly assigned to the following groups: pH1N1 (n = 15) and PRCV (n = 15). Pigs in the PRCV group were infected with PRCV/Swine/Belgium/PS-071/2020, while pigs in the pH1N1 group were infected with A/Swine/Gent/53/2019 (pH1N1), all of them intranasally. Prior to challenge, a pre-infection blood sample was drawn. Pigs from each of the PRCV and pH1N1 groups were humanely culled at 1, 5, and 12 days post-infection to assess viral load, lung pathology using Iowa and Halbur scoring, and immune responses, as described previously (14). The analysis included a total of 60 serum samples, two samples from each pig that comprised a pre-infection sample and a post-infection sample at either 1, 5 or 12 DPI, respectively. During aliquoting, a small portion of each sample was pooled and used as a quality-control (QC) sample to account for batch effects in sample processing and analysis.

For the validation phase, samples were obtained from Ghent, Belgium, following approval through the ethics committee from the Faculty of Veterinary Medicine and the Faculty of Bioscience Engineering, Ghent University (Application no:2024-041). The white Pietrain line Belgian pigs were intranasally infected with the same viruses as above: PRCV (n = 8) or pH1N1 (n = 8). Viral shedding was monitored by measuring virus titres in nasal swabs (log_10_ TCID50/100 mg) from 0-11 DPI for PRCV and 0-9 for pH1N1. Samples were collected on the day of arrival of the pigs in the experimental facilities, followed by 1, 3, 5 and 7 DPI, resulting in a total of 80 serum samples. The metadata of the samples from the discovery and validation cohort are provided in **Table S1**. A pooled validation sample was created by combining all the serum samples and was injected multiple times (n = 27) to ensure consistency throughout the analysis.

### Sample preparation and processing

Protein concentration in the serum samples was estimated by BCA assay (Thermo Fisher Scientific, UK). Serum samples equivalent to the chosen amount of protein (300 µg for discovery and 100 µg for validation studies) were aliquoted into a 96-well Protein LoBind® plate (#10215373, Eppendorf, UK). The pooled QC samples were processed alongside individual samples. The samples were randomised, and sample processing was performed using the S-TRAP™ (#P002-96MNI-0001PL, ProtiFi, NY, USA) protocol (https://protifi.com/protocols/) (31). Briefly, 10% (v/v) lysis buffer (10% sodium dodecyl sulphate (SDS), 100 mM triethylammonium bicarbonate buffer (TEAB), pH 8.5) was added to each sample containing the chosen amount of protein to achieve a final concentration of 5% (v/v) SDS, 50mM TEAB. Reduction and alkylation were carried out with a final concentration of 5 mM tris(2-carboxyethyl)phosphine (TCEP) at 55 °C for 15 minutes, followed by 20 mM iodoacetamide (IAA) at room temperature for 10 minutes in the dark. The samples were acidified with 27.5% phosphoric acid, followed by binding and washing steps with 100 mM TEAB in 90% methanol. The samples were then transferred to S-Trap™ 96-well MS sample prep kit (Protifi, USA). Proteolytic digestion was performed using Trypsin Gold (#V5280, Promega, UK) (1:20) at 47 °C for 2 hours, followed by overnight incubation at room temperature. After digestion, the peptides were eluted with 50% acetonitrile in 50 mM TEAB, then evaporated to dryness in a SpeedVac concentrator. The samples were desalted using Oasis HLB 96-well Plate, 10 mg Sorbent per Well, 30 µm (# 186000128, Waters, UK) with 80% ACN, 0.1% formic acid (FA) for column wetting, 0.1% trifluoroacetic acid (TFA) for column conditioning, sample binding and clean-up followed by 70% ACN, 0.1% FA for peptide elution. The samples were then dried down and stored at -80 °C until LC-MS/MS analysis.

For the discovery samples (n = 60), samples were reconstituted on the day of analysis in 300 µL of 98% ACN, 0.1% FA, and peptide concentration was estimated using a Nanodrop spectrometer. Indexed Retention time (iRT) (# Ki-3002-2, Biognosys, Switzerland) was added to each sample before analysis. 250 ng of peptides from each sample were injected onto the column, and the samples were analysed over 18 days, split into 9 batches. Each batch consisted of blank injections (n = 37) before and after batch but also between samples to prevent carry-over, 200 ng K562 (Promega, UK) injections (n = 4) to assess instrument performance measure instrumental variation, pooled samples injections (n = 2 batch pools, n = 1 total pool) to assess instrumental variation as well sample processing variation, and randomised swine samples (n = 9).

For the validation samples (n = 80), samples were reconstituted on the day of analysis in 250 µL 98% ACN, 0.1% FA and the peptide amount was estimated as described above. Heavy-labelled peptides synthesised and labelled at the C-terminus with a stable heavy isotope of arginine or lysine were obtained from Peptide Protein Research Ltd (n = 12) (**Table S2**) and spiked into each sample prior to analysis for retention-time confirmation and relative quantification. For each injection, 250 ng of endogenous peptides were loaded onto the LC column, spiked with 20 pg of heavy-labelled peptides, except for YYGYTGAFR (mapping to LTF), which was spiked at 100 pg. Samples were analysed in batches of 10 (n = 8) over 12 days.

### zSWATH DIA data acquisition and analysis

The samples were analysed on a ZenoTOF 7600 mass spectrometer with an Optiflow source (SCIEX, UK) coupled with an Acquity UPLC M-class liquid chromatography system (Waters, UK). Chromatographic separation was performed on a Kinetex XB-C18 LC column (2.6 µm, 0.3 mm × 150 mm, 100 Å) (# 00F-4496-AC, Phenomenex, UK) at 30 °C with a flow rate of 6 µL/min. The mobile phases A and B were water with 0.1% FA (v/v) (Optima™ LC/MS grade, #11940379, Fisher Scientific, UK) and acetonitrile (Optima™ LC/MS grade, # 10001334, Fisher Scientific, UK) with 0.1% FA, respectively. The peptides were loaded on the column using an initial 3% mobile phase B, increasing to 32% over 21 minutes, then to 80% B over 3 minutes, and held steady for 1 minute. The gradient was finally reduced to 3% B and held for 4.5 minutes to allow for column equilibration. The total gradient for this analysis was 30 minutes.

During the discovery phase, data-independent acquisition (DIA) was performed on the ZenoTOF using a Zeno SWATH acquisition scheme. The method employed 85 variable-width windows, customised based on precursor ion distributions obtained from a prior data-dependent acquisition (DDA) analysis. These windows covered a precursor mass range of 400 – 1500 m/z, and a 10 ms accumulation time was applied. Ionisation was achieved in positive mode, and the source parameters were as follows: spray voltage was set to 5000 V, curtain gas 35, CAD 7, ion source gas 1 and 2 at 20 and 60 psi respectively and a source temperature of 200 °C. Fragmentation was carried out using collision-induced dissociation (CID) in TOF MS/MS mode with a fragment mass range of 100 to 1800 *m/z*, a collision energy varying from 19 to 68 eV and an accumulation time of 13 ms.

For the validation study, a targeted mass spectrometry analysis was performed on ZenoTOF 7600 mass spectrometer, operating in multiple reaction monitoring (MRM-HR) mode with CID. A total of 24 peptides were monitored, comprising 12 endogenous target peptides and their corresponding stable isotope-labelled internal standards (IS peptides) (>97% purity). Each peptide was monitored using five-six transitions, resulting in a total of 122 MRM transitions. Transitions were defined in the TOF MS/MS settings, specifying both the precursor and fragment ion m/z value as well as the retention time for each peptide (**Table S2**). The source parameters were as follows: spray voltage was set to 5500 V, curtain gas 45, CAD 7, ion source gas 1 and 2, 20 and 60 psi respectively and a source temperature of 150 °C.

The samples were analysed in batches of 10, each batch consisting of a calibration curve prepared in a matrix and analysed alongside the samples. Calibration standards ranged from 0 to 250 pg for all peptides, except for YYGYTGAFR, for which the range was 0 to 1500 pg. In addition, pooled QC (total pool) samples were injected at regular intervals within each batch to monitor analytical performance. Blank injections were performed between all samples to minimise carryover.

### Spectral library generation

A separate gas phase fractionation LC-MS/MS method was developed for a sample-specific spectral library and applied to the pooled sample. To create a sample-specific spectral library, six individual gas-phase fractionation (GPF) LC-MS/MS methods were developed and applied to the pooled sample. They all used the same LC method, with the column and mobile phases as described in section 2.2, but with a total gradient time of 55 minutes. The gradient started with an initial 3% B, increasing to 32% over 45 minutes, then to 80% B over 2 minutes, and held steady for 2 minutes. The gradient was finally reduced back to 3% B and held for 5 min. Five of these methods used a Zeno SWATH acquisition scheme with 37 variable-width windows (vw), and the sixth used 86 vw, covering various mass ranges: 460 to 570 m/z, 570 to 680 m/z, 680 to 790 m/z, 790 to 900 m/z, and finally 900 to 1500 m/z. Other MS parameters were identical across all methods: spray voltage 5000 V, curtain gas 30, CAD 7, ion source gas 1 and 2 at 10 and 25 psi, respectively, and temperature 200 °C. The TOF MSMS parameters were the same as described above.

### Spectral library and DIA data analysis

The GPF .wiff raw files along with the sample files were used to create a project-specific library in Spectronaut (version 19.0.62635.0, Biognosys, Switzerland), which was searched against a *Sus scrofa* reference proteome database (Uniprot- UP000008227_9823, December 2024). Data obtained from DDA analysis and zSWATH (DIA) were analysed in Spectronaut version 19.0.62635.0 (Biognosys, Switzerland) using the directDIA workflow. Searches were performed against the protein database using the following parameters: peptide length of 6- 35 amino acids, Carbamidomethyl (C), set as fixed modifications, and Acetyl (protein N-term), Deamidation (N), and Oxidation (M) were set as variable modifications. Default tolerance settings were used, and 2 missed cleavages were allowed. Quantification was performed using Spectronaut’s version of MaxLFQ algorithm with precursor quantification on MS1 level. A dynamic extraction window for precursor and fragment ion detection, non-linear iRT calibration, and cross-run normalisation based on total peak area were used. All results were filtered at a q-value threshold of 0.01, corresponding to a 1% false discovery rate at both precursor and protein levels. Quantification was performed at the MS2 level using peak area integration with interference correction enabled. The list of identified proteins were exported as data matrices for further analysis.

### MRM assay development and analysis

Proteotypic peptides were selected for each candidate marker based on criteria previously described (27) as well as visual inspection of the discovery DIA data in either Spectronaut (version 19.0.62635.0) or Skyline-Daily software (24.1.1.339) (**Table S2**). Selection criteria included chromatographic peak quality and abundance, signal consistency, and the presence of multiple fragment-ion transitions supported by confident MS/MS spectra. Where suitable peptides were not observed in the discovery data (e.g. due to missed cleavages), alternative peptides were chosen based on protein sequence information. Shortlisted stable isotope-labelled internal standards (IS peptides) for 12 peptides were procured from Peptide Protein Research Ltd (>97% purity), UK.

Following peptide selection, LC-MS/MS analysis was performed to confirm fragment ion transitions and retention times. Source and compound-dependent parameters were optimised for all 12 peptides mapping to 7 proteins. Selected transitions were monitored in pooled samples containing the 12 synthetic peptides at multiple concentrations (1-1000 pg), with data acquired in technical replicates (n=3). The list of candidate proteins with their corresponding peptides selected for MRM analysis is provided in **Table S2**.

Assay performance was evaluated using calibration standards spiked into a representative serum matrix. Calibration curves were generated in technical triplicate for each calibration point using a constant background of 250 ng of endogenous peptides, with synthetic peptides ranging from 1 to 250 pg and an extended range up to 1000 pg for YYGYTGAFR mapping to LTF. Limits of detection and quantification were determined from the calibration curves, and assay reproducibility was assessed across 3 replicate measurements.

### Bioinformatics analysis

Two criteria were used for inclusion in the final analysis: >70% presence in at least one of the groups (pre-infection, pH1N1 and PRCV) and <40% CV among the total pools (n = 11). Normalisation was performed using missing value imputation 1/5 minimum positive value, group probabilistic quotient normalisation using the total pools (n = 11) and log transformed. Statistical analyses were performed in R (version 4.4.1) using packages from CRAN and the Bioconductor ecosystem, including limma (v3.62.2), ComplexHeatmap (v2.22.0), mgcv (v1.9- 3), circlize (v0.4.16), ggplot2 (v4.0.1), WGCNA (v1.73), clusterProfiler (v4.14.6), org.SS.eg.db (V3.20.0), readxl (v1.4.5), dplyr (v1.1.4), tidyr (v1.3.2), ggrepel (v0.9.6), janitor (v2.2.1), tidyverse (v2.0.0) and pROC (v1.18.0). Data export and reporting were performed using writexl (v1.5.4).

Differential protein expression analysis was performed using the limma package. Linear models with empirical Bayes moderation were fitted to log_2_-transformed protein intensities to compare post-infection time points with pre-infection. Proteins were considered significantly modulated using a threshold of |log_2_|>0.5 and Benjamini-Hochberg (BH) adjusted p-value <0.05. Weighted gene co-expression network analysis (WGCNA) was performed in R using the WGCNA package. After log_2_ transformation and variance filtering (> 0.2), modules of co-regulated proteins were identified using hierarchical clustering and dynamic tree cutting. Module eigengenes were correlated with experimental conditions to identify biologically relevant co-expression patterns.

MRM raw data were processed using SCIEX OS Analytics (V3.3.1.43), with manual verification of light- and heavy-peptide peak integration. Results were exported for downstream analysis in RStudio. Transition-level peak areas were aggregated at the peptide level using summed heavy-to-light ratios. Missing endogenous signals were imputed using a fixed low-intensity value to stabilise ratio estimation. Peptide-level ratios were log_2_- transformed prior to statistical analysis.

### Data availability

The mass spectrometry proteomics datasets supporting the conclusions of this article have been deposited in the ProteomeXchange Consortium (http://proteomecentral.proteomexchange.org) via the PRIDE partner repository, with the dataset identifier PXD076388 and PXD076418 for discovery and validation data respectively (32).

## Results

### Global analysis of proteome profiles

Longitudinal serum samples were collected from pigs challenged intranasally with either PRCV or pH1N1. Samples were taken prior to challenge (pre-infection) and at 1, 5, and 12 days post-infection (DPI) from 5 animals/group (pre-infection (n=15), 1, 5 and 12 DPI (n=5 each)) (14) (**Figure 1A, Table S1**). Viral load in nasal swabs assessed across the timepoints indicated higher nasal shedding in PRCV-infected pigs compared to those infected with pH1N1, with peak shedding occurring at 1 and 4 DPI. In both groups, viral shedding decreased and resolved by 7-8 DPI (**Figure 1B**). Lung pathology was assessed via gross and histological scoring (Halbur and Iowa scoring) as previously described (14, 33). PRCV infection resulted in more severe lung pathology at 5 DPI compared to pH1N1 (**Figures 1C-D**).

**Figure 1:**
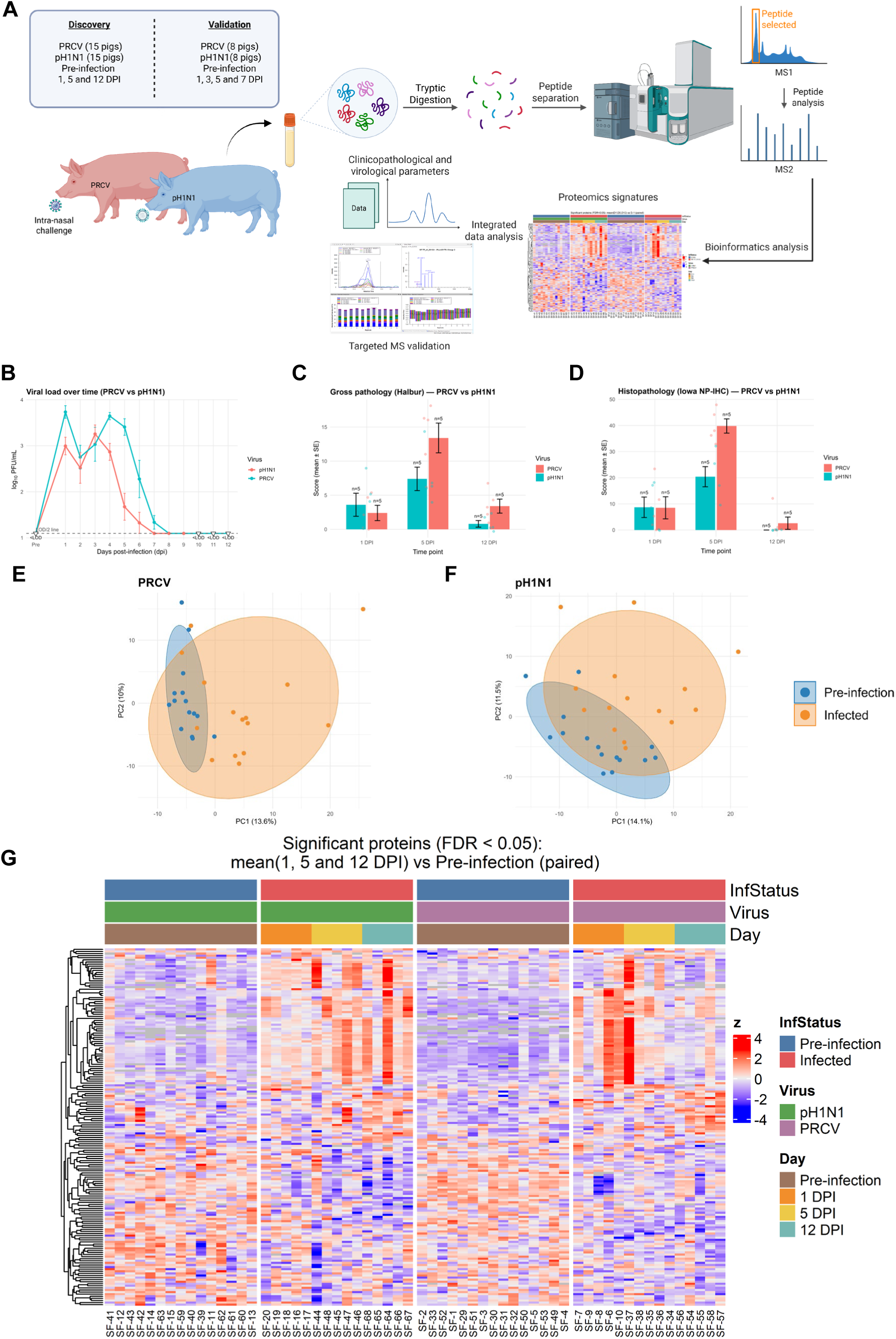
Global analysis of proteome profiles. **(A)** Schematic illustration of proteomic analysis of serum samples obtained from pigs. Illustration was created using BioRender. For the discovery phase, serum samples (n=60) were collected longitudinally from 30 pigs infected with either PRCV or pH1N1 (pre-infection; n = 15, post-infection at 1, 5 and 12 DPI; n=5 each). Candidate markers were selected based on differential analyses and validated by targeted mass spectrometry in serum samples from the Ghent cohort (N = 16 pigs, n = 80 samples). **(B)** Viral load was quantified using plaque assays from daily nasal swabs. **(C)** Gross pathology and **(D)** immunohistochemistry were used to assess lung pathology (Halbur scores), including detection of PRCV nucleocapsid (N) and pH1N1 nucleoprotein (NP) in the lungs (Iowa scores) as reported in (14). The associated clinicopathological data were integrated with the proteomics data for correlation analyses. Each point represents an individual animal; bars indicate the mean ± SEM. Statistical analysis was performed using two-way ANOVA with a Bonferroni multiple comparison correction. Principal component analysis (PCA) of serum proteomes for PRCV **(E)** and pH1N1 **(F)**, showing separation between pre-infection and post-infection samples (1, 5 and 12 DPI). **(G)** Heatmap of differentially abundant proteins comparing pre-infection with post-infection samples (PRCV in mauve and pH1N1 in green, pre-infection in blue).

Unbiased LC-MS/MS-based high-throughput proteomic analysis was performed to assess the changes in the serum proteome across the samples. To mitigate throughput bias, technical variation, and off-target depletion, STRAP-based proteomic sample preparation was applied to neat samples (31). Using a combined search of Direct DIA+ and an in-house spectral library generated from GPF and DDA analyses comprising 9,530 peptides and 1,913 unique swine proteins, we identified a total of 1,732 features corresponding to 710 protein groups. Post data processing, which included selecting proteins identified in >70% of samples within at least one group (pre-infection, pH1N1, and PRCV) and exhibiting a CV of <40% across all pools (n = 11), ∼500 unique proteins were retained for subsequent analysis. Samples were analysed across 9 batches, and following normalisation, minimal batch effects were observed (CV <15%), as evidenced by the tight clustering of the pooled samples (**Figure S1**).

We initially evaluated the global proteomic response in samples infected with pH1N1 and PRCV. The pre- and post-infection samples for both viruses demonstrated distinct clustering patterns, with pre-infection samples exhibiting tighter clustering and post-infection samples being more dispersed (**Figures 1E-F, Figures S1A-B**). Linear modelling with empirical Bayes moderation was applied to account for virus type and paired sampling within individual animals. Our analysis revealed distinct abundance patterns between pre-infection and post- infection samples. Across both infections and time points, a total of 162 proteins were identified as differentially abundant (Padj < 0.05; **Figure 1G, Figures S1C, Table S3**). Of these, 72 proteins were common to both PRCV and pH1N1, including proteins involved in the acute phase and immune response, coagulation cascade, and redox homeostasis. In pH1N1- infected samples, fifteen proteins, including PNP (F1S8H8) (log_2_FC 1.7 at 5 DPI, 1.45 at 12 DPI) and PRDX6 (Q9TSX9) (log_2_FC 1.0 at 5 DPI, 1.45 at 12 DPI), were significantly dysregulated. In contrast, 12 proteins, including APOC3 (P27917) (log2FC 0.68 at 1 DPI), TREML1 (F1RVL7) (log2FC -0.9 at 1 DPI) and CD93 (F1SAT8) (log2FC -0.6 at 1 and 5 DPI), were dysregulated in PRCV-infected samples. Notably, PRCV infection peaked as early as 1 DPI, indicating early immune response, whilst the response to pH1N1 peaked at 5 DPI and appeared more sustained, with dysregulation observed even at 12 DPI.

### PRCV- and pH1N1-specific progression signatures identify distinct early and late pathway responses

To assess temporal alterations, we next examined changes in the serum proteome at 1, 5, and 12 DPI compared to pre-infection levels in pigs challenged with PRCV and pH1N1. Principal component analysis (PCA) demonstrated distinct differences in protein expression profiles across groups. Post-infection samples (1, 5, and 12 DPI) from both PRCV and pH1N1 infections showed divergent patterns relative to pre-infection, with the most pronounced separation observed at 5 DPI (**Figures S2A-B**). Specifically, for the PRCV group, proteomics profiles from 1 DPI clustered distinctly from pre-infection, whereas the differences between pre-infection and 1 DPI for pH1N1 were relatively small, reflecting fewer proteomic differences (**Figure 1G**). Minimal differences were observed at 12 DPI in the PRCV group compared to pre-infection, with sample distributions partially overlapping pre-infection samples, in contrast to the clear segregation observed in pH1N1.

In PRCV-infected pigs, a robust early systemic response was observed at 1 DPI, characterised by 53 differentially expressed proteins compared to pH1N1 (21 proteins significantly modulated; Padj < 0.05; **Figure S2C, Table S3A**). These early PRCV-associated proteins were significantly enriched for pathways related to complement and coagulation cascades, neutrophil extracellular trap (NET) formation, carbohydrate metabolism, and hypoxia-inducible factor 1 (HIF-1) signalling (-log_10_FDR < 0.05; **Figure 2C**, **Table S3A**). This pattern is indicative of a rapid activation of innate immunity, clotting and vascular responses, and metabolic adaptation early in PRCV infection. On the contrary, in the pH1N1 group at 1 DPI, no pathways were significantly enriched, consistent with the more subtle early proteomic alterations observed in the PCA analysis (**Figure 2C, Figure S2D, Table S3A)**.

**Figure 2:**
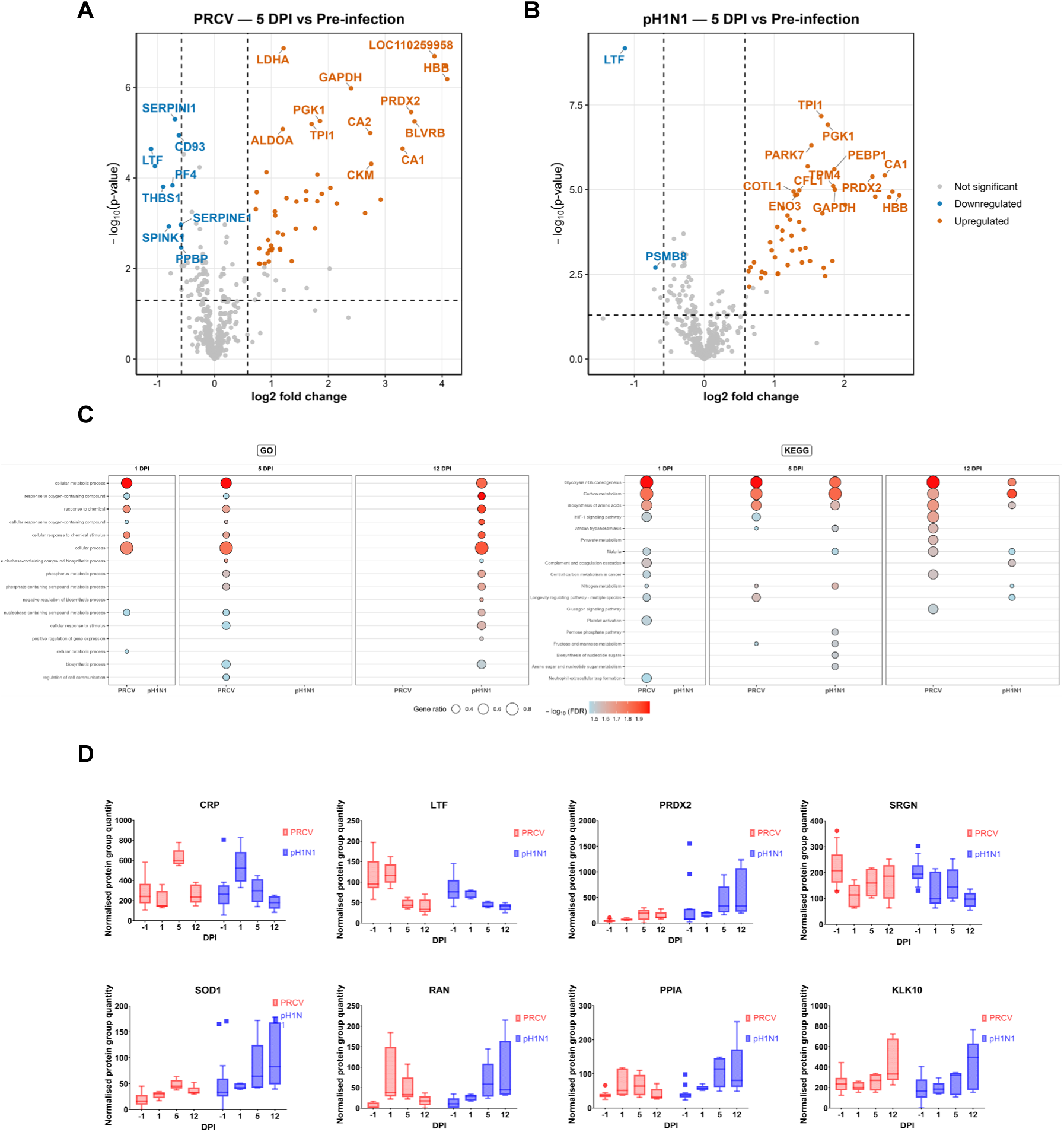
Host response signatures distinguish infection groups and reveal distinct early and late pathway responses. Volcano plots comparing serum proteome at 5 DPI in PRCV **(A)** and pH1N1 **(B)** versus pre-infection. Differential expression analysis was performed using BH- adjusted p-value. Blue and orange dots represent statistically significant proteins (adjusted p- value < 0.05 and log2FC >|0.6|) **(C)** Bubble plots representing enriched GO processes and KEGG pathways of serum proteins altered following infection with PRCV and/or pH1N1 at 1, 5, 12 DPI, based on genes list sorted by logFC values. Bubble sizes represent the ratio of the number of DE proteins associated with a specific GO processes/KEGG pathway to the total number of DE genes identified in that cluster, whilst colour represents the adjusted P-values. **(D)** Longitudinal abundance profiles for selected candidate proteins titled by gene (CRP, LTF, PRDX2, SRGN, SOD1, RAN, PPIA and KLK10) across 1,5,12 DPI for PRCV (red) and pH1N1 (blue). Boxplots illustrate the distribution of the normalised protein group intensity for each condition.

At 5 DPI, maximal proteome perturbation was observed in both virally infected groups, with broadly comparable differentially abundant proteins (pH1N1: 53 proteins; PRCV: 59 proteins; Padj < 0.05; **Figures 2A-B**). Several pathways enriched at 1 DPI in PRCV, including cellular metabolic processes, response to cellular stimulus, HIF-1 signalling, complement and coagulation, NET formation, and acute-phase response, remained significantly overrepresented at 5 DPI, indicating sustained inflammation and host defence activation. Additional pathways identified at 5 DPI in PRCV involved regulation of cell communication and biosynthetic processes. In pH1N1 group, 5 DPI marked the first time point with clear pathway-level changes, with enrichment in inflammatory response and nitrogen metabolism (- log10FDR < 0.05, **Figure 2C**). Notably, proteins involved in nucleotide sugar metabolism were enriched in pH1N1, while the longevity-regulating pathway and the HIF-1 signalling pathway were particularly enriched in PRCV infections, indicating a virus-mediated, diverse response. Thioredoxin (TXN), a key target gene associated with T-cell proliferation during Influenza A virus (IAV) infection (34), was significantly upregulated at 1 and 5DPI in PRCV compared to pH1N1, underscoring its potential role across respiratory viral infections.

By 12DPI, the PRCV group exhibited a partial return of the serum proteome towards baseline, characterised by a fewer differentially abundant proteins (25 proteins; Padj < 0.05), correlating with declining viral load (**Figure 2C, Figure S2E**). In contrast, the pH1N1 group was associated with a more persistent proteomic alteration at 12 DPI, with a higher number of altered proteins (77 proteins; Padj < 0.05) and sustained changes in inflammatory and metabolic pathways (**Figure 2C, Figure S2F**, **Table S3A**). These findings suggest that PRCV induces a strong but more transient systemic response, whereas pH1N1 elicits a more gradual yet sustained host response, consistent with the temporal clustering patterns observed in PCA. Among the proteins shared between the two datasets, Kallikrein-10 (KLK10), a secreted serine protease with anti-inflammatory and vasculoprotective properties (35), was upregulated at 12 DPI in both PRCV and pH1N1, implying a potential role in resolving or modulating late inflammatory responses. Profilin 1(PFN1), an actin-binding protein found in serum associated with preeclampsia, acute myocardial infarction, and cardiovascular disease and proinflammatory response, demonstrated sustained upregulation in pH1N1, whereas in PRCV infection, it was observed to be upregulated only on 5 DPI.

Next, building on the global analysis of PRCV- and pH1N1-specific progression signatures, we examined the longitudinal expression patterns of a subset of biologically relevant proteins that could serve as candidates for targeted validation. We focused on proteins that reflect early changes (1 DPI), persistent dysregulation (altered expression at 5 DPI and 12 DPI), and distinct trends in PRCV compared to pH1N1 (**Figure 2D**). The acute-phase proteins, C-reactive protein (CRP) and lactotransferrin (LTF), were found to differ significantly between pre- and post-infection samples. CRP levels increased in both infections, but the trends differed between PRCV and pH1N1 viral infections. While CRP levels peaked at 5 DPI in PRCV infection, levels in pH1N1 were 2-fold upregulated as early as 1 DPI, with levels returning to baseline by 5 DPI (**Figure 3C**). In the case of LTF, an iron-binding glycoprotein with varied roles in host cell defence and known antimicrobial and antiviral activity against SARS-CoV-2 and HIV (39), downregulation was observed as early as 1 DPI in pH1N1, whereas in PRCV, significant downregulation was observed only from 5 DP1. Notably, SOD1, previously identified as a serum marker of COVID-19 severity, RAN and PPIA showed peak levels at 1 DPI in the PRCV group, with levels returning to baseline by 12 DPI. Conversely, in the pH1N1 group, these proteins were significantly upregulated at 5 DPI, with a continued elevation observed even at 12 DPI, indicating a delayed but persistent response. Serglycin (SRGN), a proteoglycan, was significantly downregulated in both groups as early as 1 DPI, with PRCV showing a marked decrease compared with pH1N1. Overall, PRCV elicited a stronger and earlier systemic response than pH1N1 in agreement with other lung pathology and immunological parameters (14).

**Figure 3:**
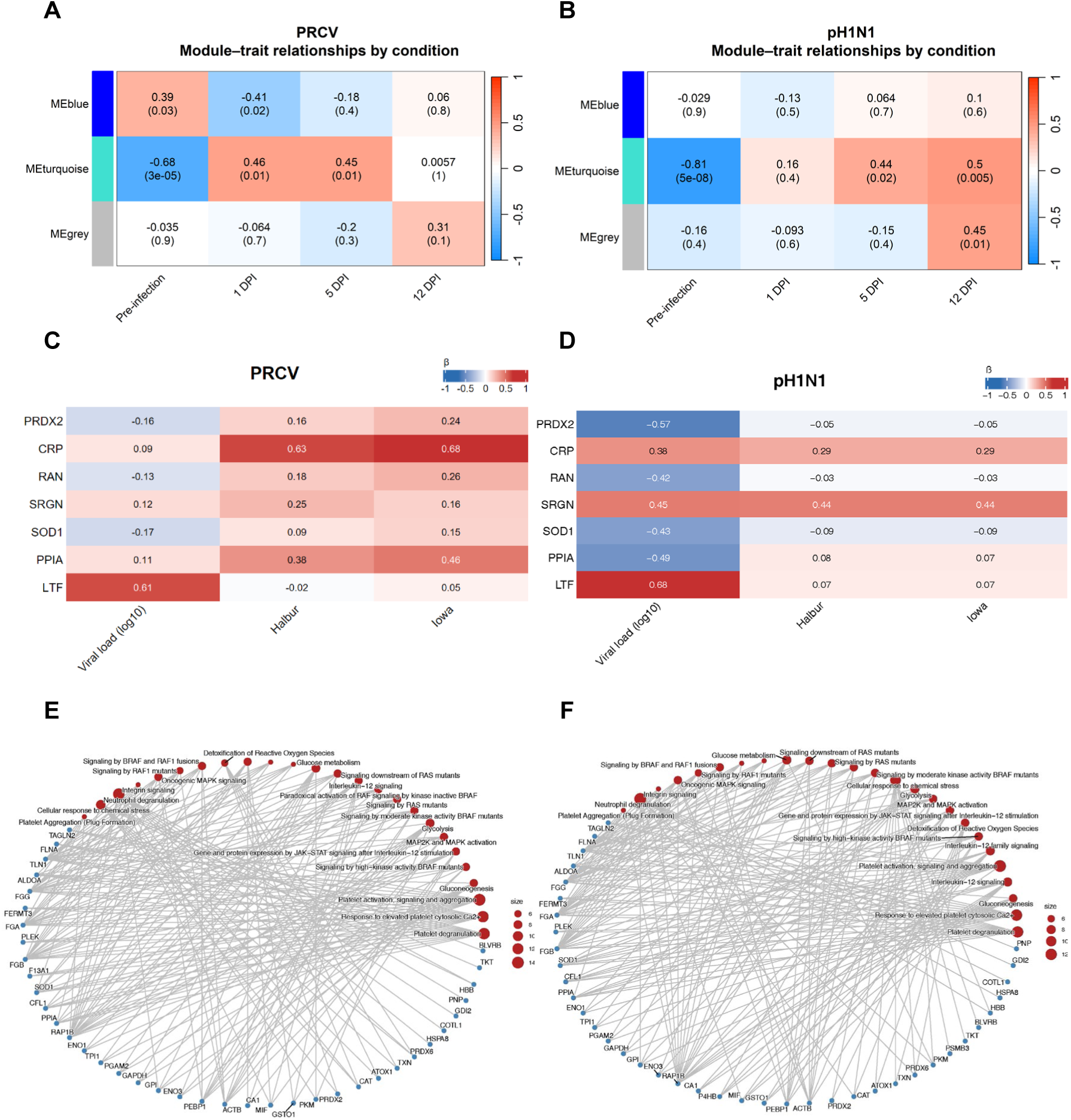
WGCNA module-trait association for PRCV **(A)** and pH1N1 **(B)**, showing correlations between module eigengenes and experimental conditions (pre-infection, 1, 5 and 12 DPI). Values indicate correlation coefficients, with adjusted P-values, where colour intensity reflects the strength and direction of the association. Protein-trait association maps for candidate features in PRCV **(C)** and pH1N1 **(D)**, showing regression coefficients (β) between protein abundance and pathology data (viral load, Halbur and Iowa scores). Positive values indicate positive association with trait severity and negative values show inverse associations. Cnetplots showing Reactome pathway enrichment of turquoise module identified from WGCNA analyses of **(E)** PRCV and **(F)** pH1N1.

### Weighted gene co-expression network analysis (WGCNA) on longitudinal samples

To further investigate co-regulated serum proteome responses and their association with the course of infection, we applied weighted gene co-expression network analysis (WGCNA) on longitudinal proteomics data for both PRCV and pH1N1 infections separately (**Figure 3A-B**). Proteins were clustered into co-expression modules, represented by their module eigengenes (blue, turquoise, and grey), and these were then correlated with infection time points (pre-infection, 1, 5 and 12 DPI).

In PRCV infection, the turquoise and blue modules exhibited the strongest associations with infection status. The turquoise module, comprising 68 proteins, showed a significant negative association with pre-infection samples (r = -0.68, P < 0.001) and significant positive correlations for post-infection at 1 and 5 DPI (r = 0.46 and 0.45, respectively, both P = 0.01). These findings suggest a low baseline abundance of these proteins prior to infection, followed by coordinated upregulation during the early and peak phases of infection, with a trend towards baseline at 12 DPI. Pathway enrichment analysis revealed that this module was significantly associated with neutrophil degranulation, reactive oxygen species (ROS) production, and interleukin-12 signalling, as well as platelet activation, signalling and aggregation (**Figure 3E**). These results are consistent with the activation of acute-phase responses, cellular stress, and innate immune activation following PRCV infection. Conversely, the blue module (n = 20 proteins) was positively associated with pre-infection samples (r = 0.39, P = 0.03) and negatively correlated with 1 DPI (r = -0.41, P = 0.02). Due to the small number of proteins, no significant pathway enrichment was observed. However, we observed several proteins mapping to immunoglobulin genes. No module showed a strong or exclusive association in 12 DPI samples, consistent with the overall proteome trending back to baseline (**Figure 3A**).

For pH1N1 infection, the turquoise module comprising 69 proteins was characterised by a negative correlation with pre-infection samples (r = -0.81, P < 0.001) followed by progressively increasing positive correlations at 5 and 12 DPI (r = 0.44 and 0.5, P = 0.02 and 0.005, respectively) (**Figure 3B**). Similar to PRCV, pathway enrichment analysis for pH1N1 revealed associations with neutrophil degranulation, reactive oxygen species (ROS) production, interleukin-12 signalling, platelet activation, signalling and aggregation (**Figure 3F**). Notably, the timing and strength of module activation differed between the two viruses. In PRCV, activation was evident as early as 1 DPI and peaked at 5 DPI, whereas in pH1N1, the increase was more gradual and remained elevated at 12 DPI, consistent with a more persistent perturbation of the serum proteome.

Overall, WGCNA demonstrated both shared (turquoise module) and virus-specific temporal patterns of serum protein co-regulation. While PRCV infection was characterised by a strong early infection response, pH1N1 infection showed a mild, progressive module response, highlighting distinct host response kinetics between these two viruses. The network-level differences corroborate pathway-level findings, supporting the conclusion that PRCV induces a rapid, intense, yet relatively transient systemic response, while pH1N1 elicits a slower yet more prolonged inflammatory and systemic response.

### Clinical associations of circulating proteins in PRCV and pH1N1 infection

We next explored associations between the differentially expressed circulating proteins over the course of infection and viral load/shedding as assessed by viral titration (PFU/mL) (**Figure 1A, Figure S3A-B**) as well as pathological parameters, including Halbur and Iowa scores (indicative of gross and histopathological severity scores). Linear regression analyses were performed between day-level mean protein abundance and clinical traits across the infection time course (**Figure S4**). The resulting regression coefficient (β) represents the direction and strength of the association between protein abundance and the clinical trait. Proteins were retained if they showed a strong association with at least one trait (|β| ≥ 0.4). Since the analysis included only the union of differentially expressed proteins, the resulting set comprises infection-responsive proteins associated with disease severity. In total, 22 proteins in PRCV and 41 in pH1N1 showed consistent associations, with 9 features shared between the viruses, suggesting a common core response to respiratory viral infection (**Table S4C-D**). These included APOC3, LTF and PRDX6; however, the directionality and dominant traits were not always conserved. APOC3 exhibited negative associations with severity traits in both infections, although only the association with pathology scores surpassed the cutoff in PRCV. LTF was positively associated with viral load in both infections, whereas PRDX6 showed contrasting behaviour, being positively associated with pathology in PRCV but negatively associated with viral load in pH1N1.

Several proteins showed virus-specific associations with pathology traits. In PRCV infection, CRP showed a strong positive association with histopathological severity, while CD44 displayed a strong negative association with pathology scores. APOC4 was negatively associated with viral load. In pH1N1 infection, several proteins were specifically associated with viral load, including PRDX2, RAN, SOD1, and PPIA, all of which showed strong negative associations. In contrast, SRGN and THBS1 exhibited a strong positive association with both viral load and pathological severity. Overall, these analyses indicate virus-specific proteomic response patterns, with PRCV infection demonstrating stronger pathology-associated signals than pH1N1.

### Targeted validation of candidate protein markers across viral infection time courses

To further validate our findings from the discovery experiment, we selected a panel of significantly dysregulated proteins for verification in an independent cohort. Serum samples were obtained from Ghent, Belgium. White Pietrain line Belgian pigs were intranasally infected with the same viruses: PRCV (n = 8) or pH1N1 (n = 8). Serum samples were collected at pre-infection (day of arrival) and at 1, 3, 5, and 7 DPI, resulting in a total of 80 samples, and were processed as described in the methods, with IS peptides spiked in prior to analysis. Candidate proteins for validation were prioritised based on several criteria including the peptide length (8–15 amino acids), proteotypicity, number of peptides identified per protein, whether the protein was a known acute phase protein, prior evidence of involvement in viral infection (including PRCV/pH1N1), localisation to extracellular or exosomal compartments, and whether peptides mapping to the swine proteome have human orthologs in the Human Plasma Peptide Atlas (https://peptideatlas.org/builds/human/plasma/) (36). Based on our analysis, 12 peptides were selected, mapping to RAN, SOD1, PPIA, PRDX2, and SRGN (two peptides each). We further chose CRP and LTF (1 peptide each), as these are known acute-phase proteins and are highly abundant, enabling assessment of analytical performance.

The analytical performance of the targeted MRM assays was evaluated using spike-in, stable heavy-isotope-labelled synthetic peptides prepared in a serum matrix. All peptides showed strong linearity (R2 > 0.98; **Figure S5**), with limits of detection (LOD) ranging from 1 – 5 pg and limits of quantification (LOQ) between 5 – 10 pg. Technical reproducibility was high (CV < 20%), demonstrating the robustness of the assays for quantitative serum measurements (**Table S5**). The quality control measures applied to the MRM assays indicated minimal variation in targeted peptide quantification across injections, as illustrated in **Figure S5.**

For the visualisation of longitudinal trends, protein abundances obtained from discovery DIA and validation MRM were centred on the pre-infection baseline (median abundance), calculated separately for each cohort, virus, and protein. Centred values were calculated by subtracting this baseline from each sample on the log_2_ scale. Overall, the discovery and validation datasets revealed concordance for two candidate proteins (SRGN and LTF), which exhibited broadly similar longitudinal trends across cohorts. Peptides mapping to four proteins, including PRDX2, PPIA, RAN, and SOD1, showed delayed or discordant responses. The longitudinal trend for CRP was discordant for both viruses (**Figure 4A-D, Figure S6**). These observed results could reflect differences between studies, including pig breed, country of origin, management conditions and sampling schedules.

**Figure 4:**
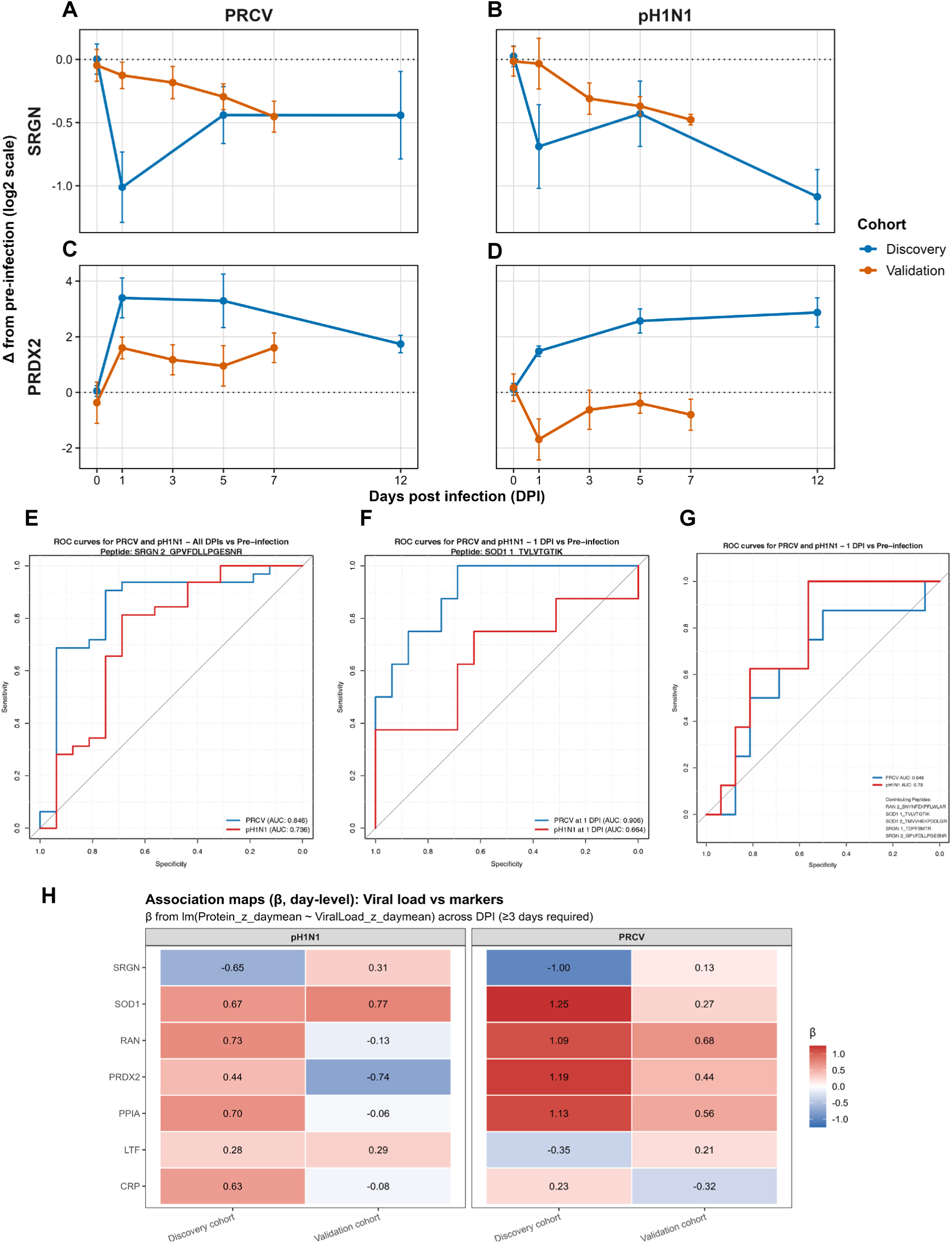
Centred time-course profiles of serum protein levels for SRGN **(A – B)** and PRDX2 **(C – D)** in PRCV **(A, C)** and pH1N1 **(B, D)** infection. Protein abundances from the discovery and validation (MRM) cohorts were centred on the median pre-infection baseline, separately for each cohort, virus, and protein. Values represent mean log_2_ change from pre-infection ± standard error across DPI. **(E)** ROC curves for SRGN peptide discriminating infected versus pre-infection samples across all DPIs in PRCV (AUC 0.846) and pH1N1 (AUC 0.736). **(F)** ROC curves for SOD1 peptide discriminating 1 DPI versus pre-infection in PRCV (AUC 0.906) and pH1N1 (AUC 0.664). **(G)** ROC curves for combined peptide panel (RAN, SRGN and SOD1) discriminating 1 DPI versus pre-infection in PRCV (AUC 0.648) and pH1N1 (AUC 0.75). **(H)** Viral load association map comparing candidate markers in the discovery and validation cohort. Associations represent regression coefficients from day-level linear models describing the relationship between protein abundance and viral load across all time points for PRCV and pH1N1.

For pH1N1 infection, several proteins exhibited comparable patterns across cohorts, particularly at early to mid-stages of the infection. SRGN displayed closely correlated longitudinal profiles across all days post-infection, indicating consistent regulation across cohorts. SOD1 also showed strong concordance at early time points, with a notable increase at 1 DPI in both datasets. Other candidates, including LTF, showed a milder agreement between cohorts, with limited changes in abundance over time. Whilst the overall direction of change was comparable, the magnitude of regulation differed between discovery and validation datasets. Several proteins displayed discordant temporal patterns across cohorts. CRP showed a pronounced early increase in the discovery cohort, followed by a return to baseline, whereas the validation cohort exhibited a marked early decrease post-infection. PPIA demonstrated opposite trends and PRDX2 did not show evidence of upregulation in the validation cohort, remaining below baseline relative to pre-infection samples across the time course.

For PRCV infection, a similar pattern was observed between the two cohorts. SRGN showed strong alignment across cohorts throughout the time course, indicating consistent regulation. PRDX2 and RAN also demonstrated good agreement between discovery and validation datasets, supporting their reproducibility in the PRCV model. Other candidates, such as LTF and PPIA, showed only mild concordance, resulting in mild agreement between cohorts. In contrast, CRP and SOD1 displayed opposing trends between cohorts (**Figure 4A-D**).

To evaluate the diagnostic potential of candidate markers, receiver operating characteristic (ROC) analyses were conducted on peptide-level abundances (heavy-to-light ratios) derived from the validation dataset. Discrimination between pre- and post-infection samples was assessed both across all DPI combined and individually at 1 DPI, aiming to investigate early detection (**Figure S7**).

Across all DPI (**Figure 4E**), several peptides demonstrated moderate to strong discriminatory performance. The SRGN-derived peptide GPVFDLLPGESNR exhibited consistent accuracy across both PRCV and pH1N1, with the respective areas under the curve (AUC) of 0.846 and AUC 0.736. The peptide TVLVTGTIK, mapping to SOD1, showed the strongest performance, with AUCs of 0.945 and 0.879 for PRCV and pH1N1, respectively. Additionally, RAN peptide SNYNFEKPFLWLAR displayed good discrimination (PRCV AUC 0.777, pH1n1 AUC 0.719), whereas PRDX2, PPIA and LTF peptides generally exhibited limited diagnostic utility.

Focusing on early infection (1 DPI vs pre-infection, **Figure 4F**), SOD1 remained the most effective early marker for PRCV (AUC 0.906), while SRGN and RAN peptides showed moderate but reproducible performance across both infections. A combined multi-peptide panel including SOD1, SRGN and RAN showed moderate early classification performance (PRCV AUC 0.65, pH1N1 AUC 0.75, **Figure 4G**). Although these combined markers did not outperform the strongest individual biomarkers, they help to capture complementary host-response pathways and may thus enhance the robustness of the classification.

Overall, the comparison of PRCV and pH1N1 revealed differences in longitudinal marker behaviour across infection models. SRGN demonstrated consistent agreement between discovery and validation cohorts for both viruses. RAN and PRDX2 displayed similar trends exclusively in PRCV infection. SOD1 showed a concordant and reproducible increase during the early stages of pH1N1 infection (1 DPI), although it remained below baseline levels in the PRCV validation cohort. LTF and PPIA displayed very mild changes with partial concordance across cohorts and viruses. CRP, by contrast, showed opposite trends between the discovery and validation cohorts for both viruses (**Figure S6**).

### Association of candidate protein markers with viral load and lung pathology in the validation cohort

We next assessed the longitudinal associations between candidate protein abundance and viral load (log_10_ TCID50 per 100 mg of tissue) across all time points (**Figure 4H**). For each protein, linear models were fitted separately within PRCV and pH1N1 infection groups, with regression coefficients (β) used to quantify the direction and magnitude of association.

Day-level association analyses revealed distinct discovery-to-validation patterns for both viruses. In PRCV infection, SOD1, RAN, PRDX2, and PPIA demonstrated consistently positive associations with viral load in both the discovery and validation datasets, although effect sizes were smaller in the validation cohort. In the pH1N1 group, SOD1 consistently showed positive associations across both cohorts, whereas LTF exhibited a weak positive association throughout. In contrast, several markers revealed discordant patterns between discovery and validation. In PRCV, SRGN exhibited a strong negative association in the discovery study but a mild positive association in the validation study. LTF shifted from a mild negative to a mild positive association, whereas CRP showed the opposite trend. Greater discrepancies were observed in pH1N1 infection, with RAN, PPIA and CRP showing strong positive associations with viral load in discovery but mild negative associations in the validation study. Notably, PRDX2 demonstrated the largest change, shifting from a positive association in discovery to a strong negative association in validation. SRGN also changed from a negative to a mild positive association between cohorts.

## Discussion

This study investigated longitudinal changes in swine following coronavirus or influenza infections using a discovery approach with zSWATH-MS, followed by targeted validation employing an MRM-HR method. The analysis revealed changes in protein expression following infection with the respiratory viruses PRCV and pH1N1, highlighting virus-specific host responses. Multivariate analyses showed that while pre-infection samples remained tightly clustered, post-infection samples were more dispersed, indicating infection-induced biological variation. Even though the infection-specific PCAs did not reveal a strong separation between pre-infection and infected samples, they showed consistent clustering of the former and progressive shifts in infected samples over time, with 1 and 5 DPI being the most distinct from pre-infection samples in the case of PRCV infection. These results support the need for a proteomic profiling approach to identify early systemic responses to infection that correlate with increased viral load early during infection and continue to remain affected when pathological changes within the lungs are evident at 5 DPI.

Longitudinal data further revealed temporal changes in several proteins, with both common and virus-specific trends. The temporal separation observed in PCA is consistent with the time-resolved differential expression and WGCNA modules. C-reactive protein and lactotransferrin, two known markers of infection in humans (37, 38) were significantly altered post-infection in the discovery cohort. For both PRCV and pH1N1, we see upregulation of CRP protein and downregulation of LTF. However, the CRP trend differs between conditions, with protein levels peaking earlier in pH1N1 (1 DPI) than in PRCV (5 DPI). This suggests potential differences in the timing of the acute-phase response between PRCV and pH1N1 viruses.

In addition to these known markers, several proteins emerged as potential candidates for further investigation during the discovery phase. These include: PRDX2, involved in oxidative stress and inflammation; Superoxide Dismutase 1 (SOD1), an antioxidant enzyme; Peptidylprolyl isomerase A (PPIA), implicated in immune modulation; RAN, a GTP-binding protein with nuclear transport function; and Serglycin (SRGN), a secretory vesicle proteoglycan associated with immune cell granules (39).

Peroxiredoxin 2 (PRDX2) is a thiol-specific antioxidant protein belonging to the Peroxiredoxin family, which is known to counter oxidative stress induced by pathogens, including bacteria (40) and viruses (41, 42). PRDX2 was found to be secreted in the supernatant of A549 lung epithelial cells in response to influenza infection (41).

SOD1 is a well-characterised antioxidant enzyme responsible for scavenging free radicals in cells. Increased SOD1 expression has been observed in patients with asymptomatic influenza A infection, whereas in vitro studies have shown that H5N1 and RSV infections in lung epithelial cells lead to decreased SOD1 expression, resulting in significant oxidative stress and increased reactive oxygen species (ROS) production that facilitates viral replication and inflammation (43–45). Serum SOD1 levels have recently been identified as a potential predictive marker of SARS-CoV-2 progression, indicating COVID-19 severity. Results from our targeted validation further corroborate these findings, demonstrating SOD1 as the most effective early marker for PRCV (AUC 0.906). Together, SOD1 could serve as a potential marker for assessing zoonotic spillover. Additionally, the same study also reports increased PRDX2 and LDHA levels in patients with severe symptoms, supporting our findings from the PRCV group. SOD1 has been reported to be secreted extracellularly via unconventional pathways in nasopharyngeal cell exosomes (47), platelet exosomes (48) and through Sec61 translocon in *Saccharomyces cerevisiae* (49).

Peptidylprolyl isomerase A (PPIA), also known as Cyclophilin A is an enzyme that catalyses the cis-trans isomerisation of proline imidic peptide bonds, especially in proteins with intrinsically disordered regions (IDRs) (50), a key factor that accelerates protein folding (51). A previous study by Gamble and colleagues showed that PPIA can interact with the HIV-1 (human immunodeficiency virus type 1) gag protein (52), enhance viral infection, and that PPIA inhibitors decrease HIV-1 replication (53). Further, Liu and colleagues showed that PPIA interacts with the M1 protein of Influenza A virus and impairs viral replication (54). Similarly, PPIA has been suggested to be involved in the replication of hepatitis C virus and human coronaviruses (55, 56). These findings suggest that PPIA could also be important in pH1N1 and PRCV infection. RAN, a Ras-related nuclear (RAN) protein, is a small GTP-binding protein that is involved in the translocation of RNA and proteins through the nuclear pore complex. It has been previously known to be important for the viral replication of Hepatitis C Virus (57) and Influenza A Virus (58).

The candidate markers- PRDX2, SOD1, PPIA, RAN and SRGN were chosen for validation based on their novelty and their biological relevance, particularly in host immune and stress response pathways. The validation experiments using targeted MRM assays were performed to assess whether the temporal protein changes identified in the discovery study could be reproduced in an independent cohort. Overall, agreement between discovery and validation was limited and varied across candidate protein markers and viruses. Some candidates showed consistent behaviour across studies, whilst others displayed attenuated, delayed or opposite trends in the validation data. Across both PRCV and pH1N1 infections, SRGN showed the most consistent behaviour with similar longitudinal trends observed in both discovery and validation datasets. This was also reflected in the viral load analyses, where SRGN showed a consistent negative association with pH1N1. These findings indicate that SRGN regulation is relatively stable across infection models and analytical approaches. Interestingly, SRGN transcript levels were found to be significantly dysregulated in mice lungs from re-challenge groups infected with IAV-X31 compared to primary infection (59).

PRDX2 showed a mixed response; the longitudinal effect was reproduced in the PRCV model but not under pH1N1 infection. It also showed partial agreement with viral load and pathology, particularly for pH1N1 viral load and PRCV pathology scores. SOD1 showed a clear early post-infection response to pH1N1 that was reproduced in the validation study, but this behaviour was not observed for PRCV. The rest of the candidates, PPIA, RAN, LTF, and CRP, showed weaker or discordant trends across cohorts, depending on the virus.

Several factors may explain why some candidate proteins identified in the discovery study did not validate consistently. Firstly, differences in pig breed, origin, and experimental setup between the discovery and validation cohorts could influence baseline immune status and the host response to infection. Secondly, protein-level trends were more consistent across time in the discovery dataset than in the validation dataset. Although early sampling time points were comparable between cohorts, variations in sampling methods may have contributed to the discrepancies observed, as discovery samples were collected postmortem, whereas validation samples were obtained longitudinally from live animals. Nonetheless, several markers tested with the targeted approach showed largely similar trends. Variability in pig breeds, origins, immune heterogeneity, sampling conditions, and the use of samples from live versus deceased animals reflect the inherent complexities of field sampling in real-world settings, especially in a pandemic scenario. Any diagnostic test developed from work such as this should show consistent trends across breeds to ensure robustness. But, methodological differences between SWATH DIA-based discovery and targeted MRM assays may also contribute, as the former reflects protein-level changes inferred from multiple peptides, whereas targeted MRM assays rely on a small number of predefined peptides that may not fully capture the overall protein behaviour.

The results from the discovery and validation studies suggest that validating candidate protein markers for respiratory infections is complex. Instead of identifying universally reproducible markers, the findings point towards virus-specific and time-dependent protein responses. Here, SRGN has emerged as a promising candidate marker. It showed consistent longitudinal response across cohorts and viruses, with concordance with viral load and pathology data. This suggests that SRGN may represent more stable components of the host response, whereas others may be informative only within specific infections or time windows, or when interpreted alongside viral load and pathology scores.

In addition to longitudinal reproducibility, ROC analyses provided further insight into the diagnostic potential of selected candidates. SRGN showed consistent discriminatory performance across infection models and time points, reinforcing its stable regulation observed between discovery and validation cohorts. SOD1 demonstrated strong classification performance at early stages of infection, particularly at 1 DPI, suggesting potential utility as an early infection marker. Although individual peptides showed variable performance across PRCV and pH1N1, combining markers may improve classification robustness across infection models and reduce reliance on single-peptide measurements. These findings remain exploratory, as diagnostic performance was assessed in a limited validation cohort. Further evaluation in larger and more diverse populations will be required to determine the translational utility of these candidate biomarkers.

## Conclusions

Taken together, our findings demonstrate that longitudinal serum proteomics in a natural porcine host model provides a powerful framework for dissecting host responses to respiratory virus infection, directly relevant to veterinary and human disease. By comparing temporal proteomic trajectories following influenza and PRCV infection, we identified both shared and virus-specific patterns associated with disease severity, thereby revealing mechanistic differences in pathogenesis that are not apparent from clinical signs or virological measurements alone. Furthermore, the development of targeted assays and the validation of four candidate proteins, such as SRGN, PRDX2, SOD1, and RAN, which correlate with viral load and pathological features, emphasise the potential usefulness of these assays for differentiating respiratory viral aetiologies and categorising disease severity in veterinary contexts. Overall, our findings provide a resource for prioritising host-response markers and pathways that may be translatable to human influenza and coronavirus infections.

## List of abbreviations

pH1N1: Swine influenza A virus
PRCV: Porcine respiratory coronavirus
CoV: Coronaviruses
DPI: days post-infection
AUC: Area under the curve
SRGN: Serglycin
SOD1: Superoxide dismutase 1
LC-MS/MS: Liquid chromatography mass spectrometry
BCA: Bicinchoninic acid
QC: Quality control
TEAB: Triethylammonium bicarbonate
FA: Formic acid
ACN: Acetonitrile
TFA: Trifluoroacetic acid
iRT: Indexed Retention time
DIA: Data-independent acquisition
DDA: Data-dependent acquisition
CID: Collision-induced dissociation
GPF: Gas phase fractionation
MRM-HR: Multiple reaction monitoring
WGCNA: Weighted gene co-expression network analysis
CV: coefficient of variation
PCA: Principal component analysis
ROC: Receiver operating characteristics

## Declarations

## Ethics approval and consent to participate

Serum samples for the discovery phase were collected from an infectious study at the Pirbright Institute, UK, following approval by the ethics review process under the Animals (Scientific Procedures) Act 1986, with project licences PP7764821 and PP2064443. The study received a favourable retrospective governance review (FRGR) from the University of Surrey NASPA, a sub-committee of the Animal Welfare and Ethical Review Board (NASPA-2526-08(L)).

For the validation phase, samples were obtained from Ghent, Belgium, following approval through the ethics committee from the Faculty of Veterinary Medicine and the Faculty of Bioscience Engineering, Ghent University (Application no:2024-041).

## Consent for publication

Not applicable

## Author’s Contributions

C.F. designed and performed experiments, analysed data, and wrote the original manuscript. B.P. and J.G performed experiments and revised the manuscript. A.D.W. and K.v.R. contributed to the resources and to the manuscript. E.T., Y.S. and S.M.P. acquired funding, wrote and revised the manuscript. S.M.P. conceived the study, designed experiments, supervised, and administered the research. All authors were involved in reviewing and editing.

## Acknowledgments

We are grateful to the animal staff at The Pirbright Institute for providing excellent animal care. We thank Victoria South at SCIEX for their support and technical expertise in developing the MRM method on the ZenoTOF 7600 platform.

## Additional files

**Additional file 1 contains Figures S1 – S7**

**Additional file 2 contains Tables S1 – S6**

