## Supplementary Figures for "Longitudinal serum proteomics analyses reveal early diagnostic and discriminatory markers for porcine influenza and coronavirus infections": Supplemental files.pdf

**Figure legends**

**Figure S1:** PCA plots of pre-infection, pH1N1, and PRCV-infected serum samples analysed across nine batches demonstrate minimal segregation in the pre-infection control samples **(A)**. Post-normalisation, minimal batch effects were observed, evidenced by the tight clustering of the total pools **(B)**. **(C)** Venn diagram depicting the number of unique and shared differentially regulated proteins (Padj < 0.05) between PRCV-infected pigs and pH1N1-infected pigs.

**Figure S2:** PCA plots reveal differences in protein expression across groups, with post-infection samples (1, 5, and 12 DPI) for both PRCV **(A)** and pH1N1 **(B)** infections exhibiting distinct patterns compared to pre-infection samples, largely driven by the 5 DPI samples. Volcano plots comparing serum proteome in PRCV and pH1N1 versus pre-infection at 1 DPI **(C-D)** and 12 DPI **(E-F)**. Differential expression analysis was performed using linear modelling with empirical Bayes moderation in the *limma* R package. Orange and blue dots represent statistically significant proteins (adjusted p-value < 0.05 and log2FC >|0.6|).

**Figure S3:** Protein-trait association maps for all differentially expressed significant proteins in PRCV **(A)** and pH1N1 **(B)**, showing regression coefficients (β) between protein abundance, viral load and pathology data (Halbur and Iowa scores).

**Figure S4:** Viral load correlation for all differentially expressed significant proteins in PRCV **(A)** and pH1N1 **(B)** across all time points. Heatmaps show standardised protein abundance (Z-score, day mean).

**Figure S5:** Evaluation of analytical performance of selected peptides for targeted MRM assay development. **(A-L)** Linearity curve to determine the limits of detection and quantitation of selected peptides mapping to CRP, LTF, PPIA, PRDX2, RAN, SOD1 and SRGN. A dilution series of the peptides was carried out by spiking a pooled set of 12 isotopically labelled synthetic peptides into a pooled serum matrix at different concentrations (5–500 fmol). Linear regression analysis was performed using Analyst software, with a maximum LOQ bias of xx and an LOQ CV of 20%. Analyses were performed in triplicate on a Sciex ZenoToF 7600 mass spectrometer using MRM-HR mode. Error bars represent SD for triplicates. All peptides showed strong linearity (R2 > 0.98), with limits of detection of 1 – 5 pg and limits of quantification of 5 – 10 pg. **(M)** Log2-transformed peptide abundances for the 12 monitored peptides plotted across injection order before (Pre, red) and after (Post, blue) total pool normalisation. Each panel represents a single peptide and illustrates improved signal stability across injections after normalisation.

**Figure S6:** Centred time-course profiles of serum protein abundance for all 7 candidate markers for PRCV (**A**) and pH1N1 (**B**). Protein abundances from the discovery (DIA-MS) and validation (MRM) cohorts were centred on the median pre-infection baseline, separately for each cohort, virus, and protein. Values represent mean $\log_2$ change from pre-infection ± standard error across days post-infection. **(C)** Viral load kinetics in the validation cohort were quantified and expressed as log10 TCID50 per 100 mg of tissue over the infection time course for PRCV and pH1N1.

**Figure S7:** Receiver operating characteristic (ROC) curves for individual peptides discriminating infected versus pre-infection samples in PRCV and pH1N1 across all post-infection timepoints (1, 5 and 12 DPI; **A – K**) and restricted to 1 DPI (**L – V**). Each panel corresponds to a single peptide mapping to one of the candidate proteins (CRP, LTF, PPIA, SRGN, SOD1, RAN or PRDX2). Discriminatory performance is reported as the area under the curve (AUC) for both PRCV and pH1N1.

**Figure S1**

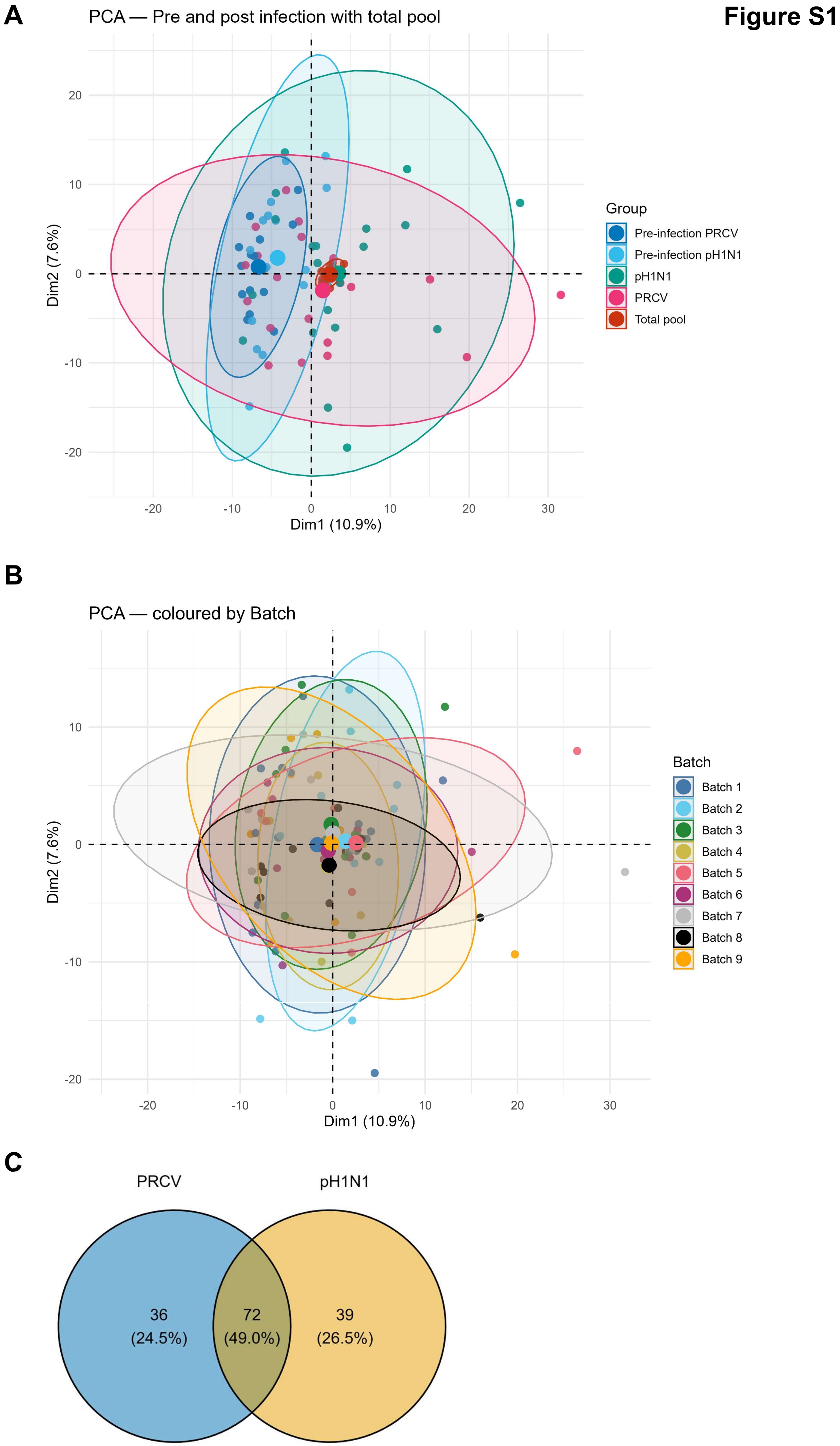
A
PCA — Pre and post infection with total pool
Dim1 (10.9%)
Dim2 (7.6%)
Group
Pre-infection PRCV
Pre-infection pH1N1
pH1N1
PRCV
Total pool
B
PCA — coloured by Batch
Dim1 (10.9%)
Dim2 (7.6%)
Batch
Batch 1
Batch 2
Batch 3
Batch 4
Batch 5
Batch 6
Batch 7
Batch 8
Batch 9
C
PRCV
pH1N1
36
(24.5%)
72
(49.0%)
39
(26.5%)

**Figure S2**

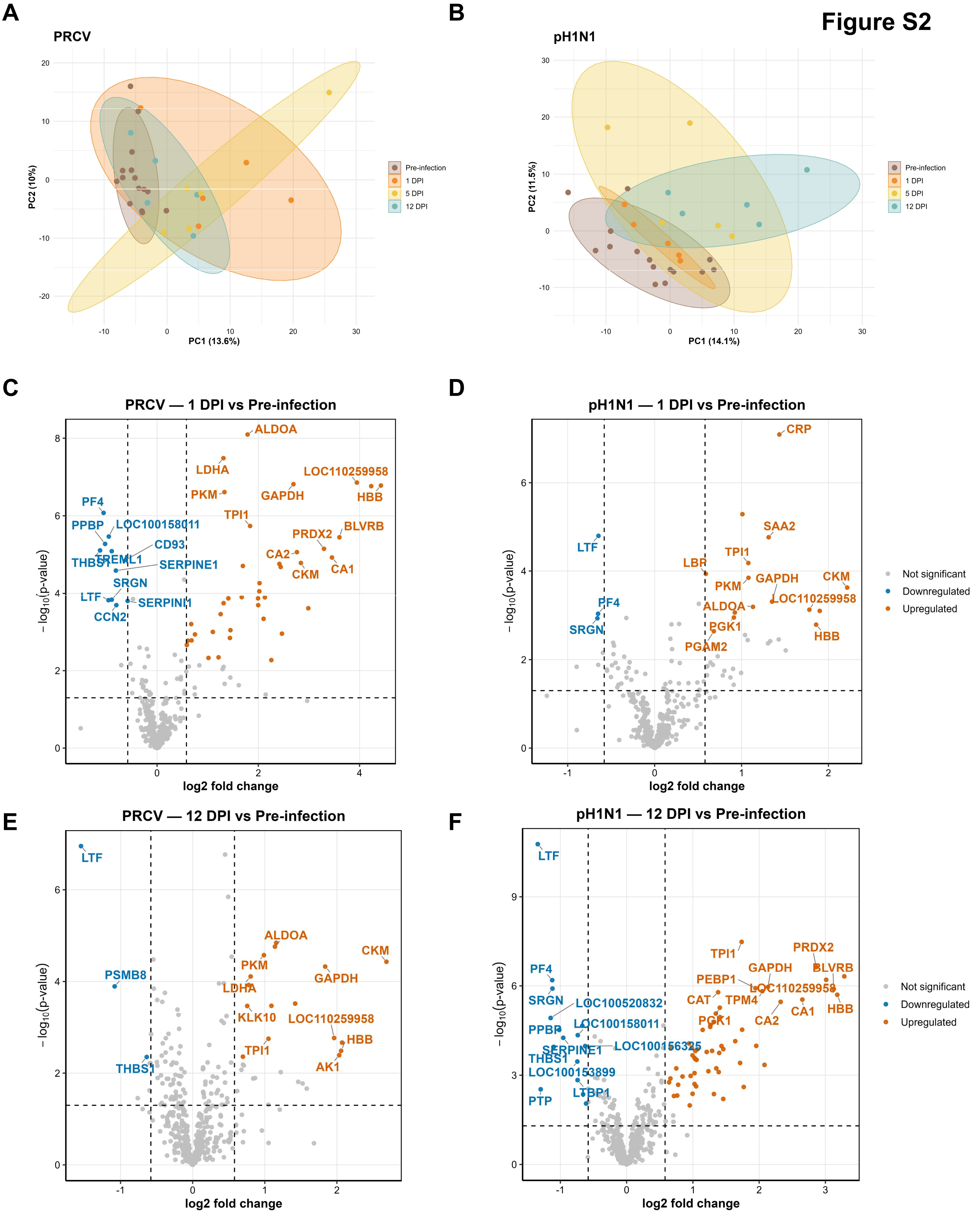
A
PRCV
PC1 (13.6%)
PC2 (10%)
Pre-infection
1 DPI
5 DPI
12 DPI
B
pH1N1
PC1 (14.1%)
PC2 (11.5%)
C
PRCV — 1 DPI vs Pre-infection
ALDOA
LDHA
PKM
GAPDH
LOC110259958
HBB
TPI1
PRDX2
BLVRB
CA2
CKM
CA1
PF4
PPBP
LOC100158011
CD93
THBS1
TREML1
SERPINE1
SRGN
LTF
SERPINI1
CCN2
log2 fold change
−log10(p-value)
D
pH1N1 — 1 DPI vs Pre-infection
CRP
SAA2
LTF
TPI1
LBP
PKM
GAPDH
CKM
LOC110259958
ALDOA
PF4
SRGN
PGK1
PGAM2
HBB
Not significant
Downregulated
Upregulated
E
PRCV — 12 DPI vs Pre-infection
LTF
ALDOA
CKM
PKM
PSMB8
LDHA
GAPDH
KLK10
LOC110259958
TPI1
HBB
THBS1
AK1
F
pH1N1 — 12 DPI vs Pre-infection
LTF
PRDX2
TPI1
GAPDH
BLVRB
PF4
PEBP1
LOC110259958
SRGN
LOC100520832
CAT
TPM4
CA1
HBB
CA2
PPBP
LOC100158011
PGK1
SERPINE1
LOC100156325
THBS1
LOC100153899
LTBP1
PTP
Not significant
Downregulated
Upregulated

#### Figure S3

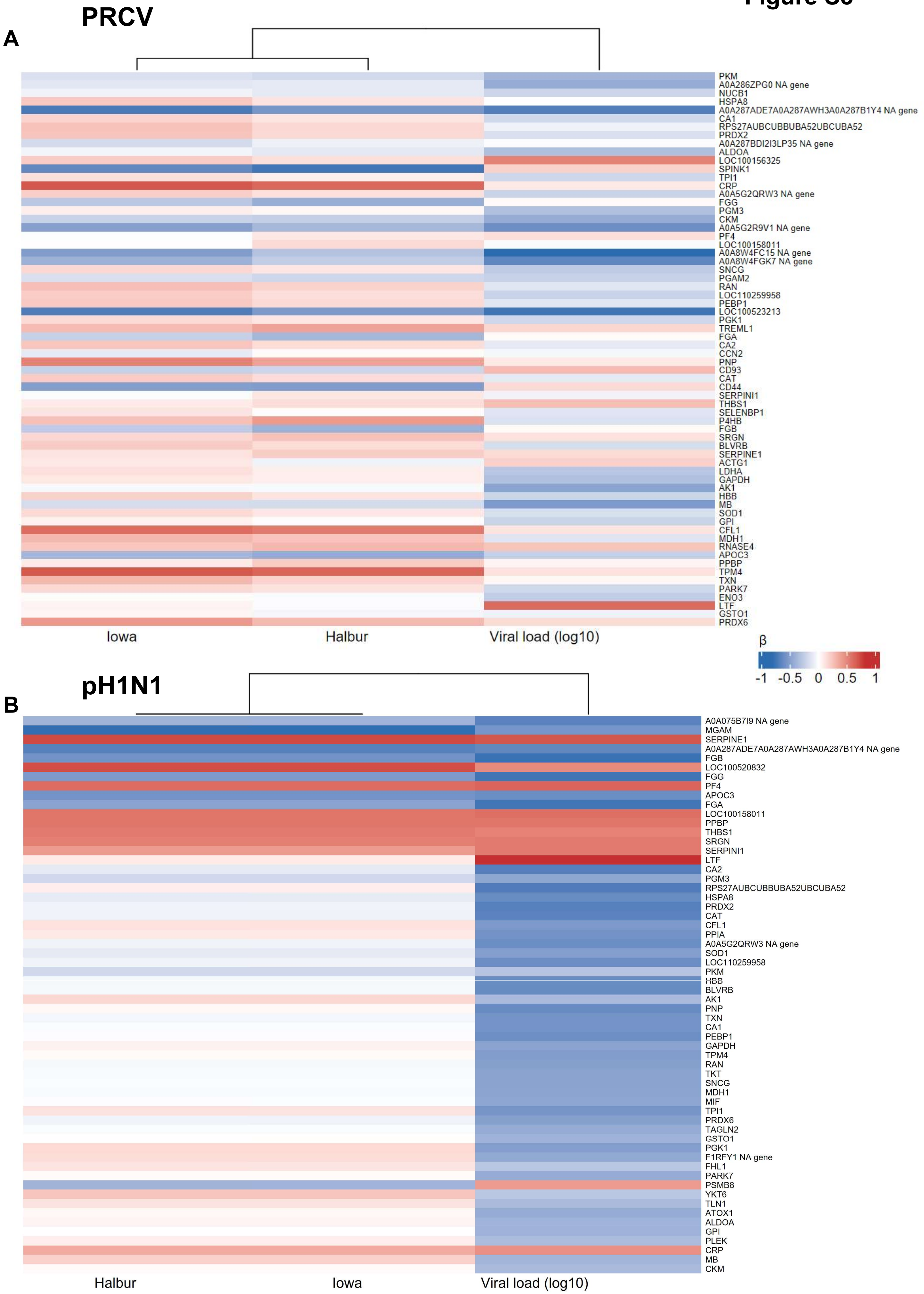
PRCV
A
PKM
A0A286ZPG0 NA gene
NUCB1
HSPA8
A0A287ADE7A0A287AWH3A0A287B1Y4 NA gene
CA1
RPS27AUBCUBBUBA52UBCUBA52
PRDX2
A0A287BDI2I3LP35 NA gene
ALDOA
LOC100156325
SPINK1
TPI1
CRP
A0A5G2QRW3 NA gene
FGG
PGM3
CKM
A0A5G2R9V1 NA gene
PF4
LOC100158011
A0A8W4FC15 NA gene
A0A8W4FGK7 NA gene
SNCG
PGAM2
RAN
LOC110259958
PEBP1
LOC100523213
PGK1
TREML1
FGA
CA2
CCN2
PNP
CD93
CAT
CD44
SERPINI1
THBS1
SELENBP1
P4HB
FGB
SRGN
BLVRB
SERPINE1
ACTG1
LDHA
GAPDH
AK1
HBB
MB
SOD1
GPI
CFL1
MDH1
RNASE4
APOC3
PPBP
TPM4
TXN
PARK7
ENO3
LTF
GSTO1
PRDX6
Iowa
Halbur
Viral load (log10)
β
-1
-0.5
0
0.5
1
pH1N1
B
A0A075B7I9 NA gene
MGAM
SERPINE1
A0A287ADE7A0A287AWH3A0A287B1Y4 NA gene
FGB
LOC100520832
FGG
PF4
APOC3
FGA
LOC100158011
PPBP
THBS1
SRGN
SERPINI1
LTF
CA2
PGM3
RPS27AUBCUBBUBA52UBCUBA52
HSPA8
PRDX2
CAT
CFL1
PPIA
A0A5G2QRW3 NA gene
SOD1
LOC110259958
PKM
HBB
BLVRB
AK1
PNP
TXN
CA1
PEBP1
GAPDH
TPM4
RAN
TKT
SNCG
MDH1
MIF
TPI1
PRDX6
TAGLN2
GSTO1
PGK1
F1RFY1 NA gene
FHL1
PARK7
PSMB8
YKT6
TLN1
ATOX1
ALDOA
GPI
PLEK
CRP
MB
CKM
Halbur
Iowa
Viral load (log10)

**Figure S4**

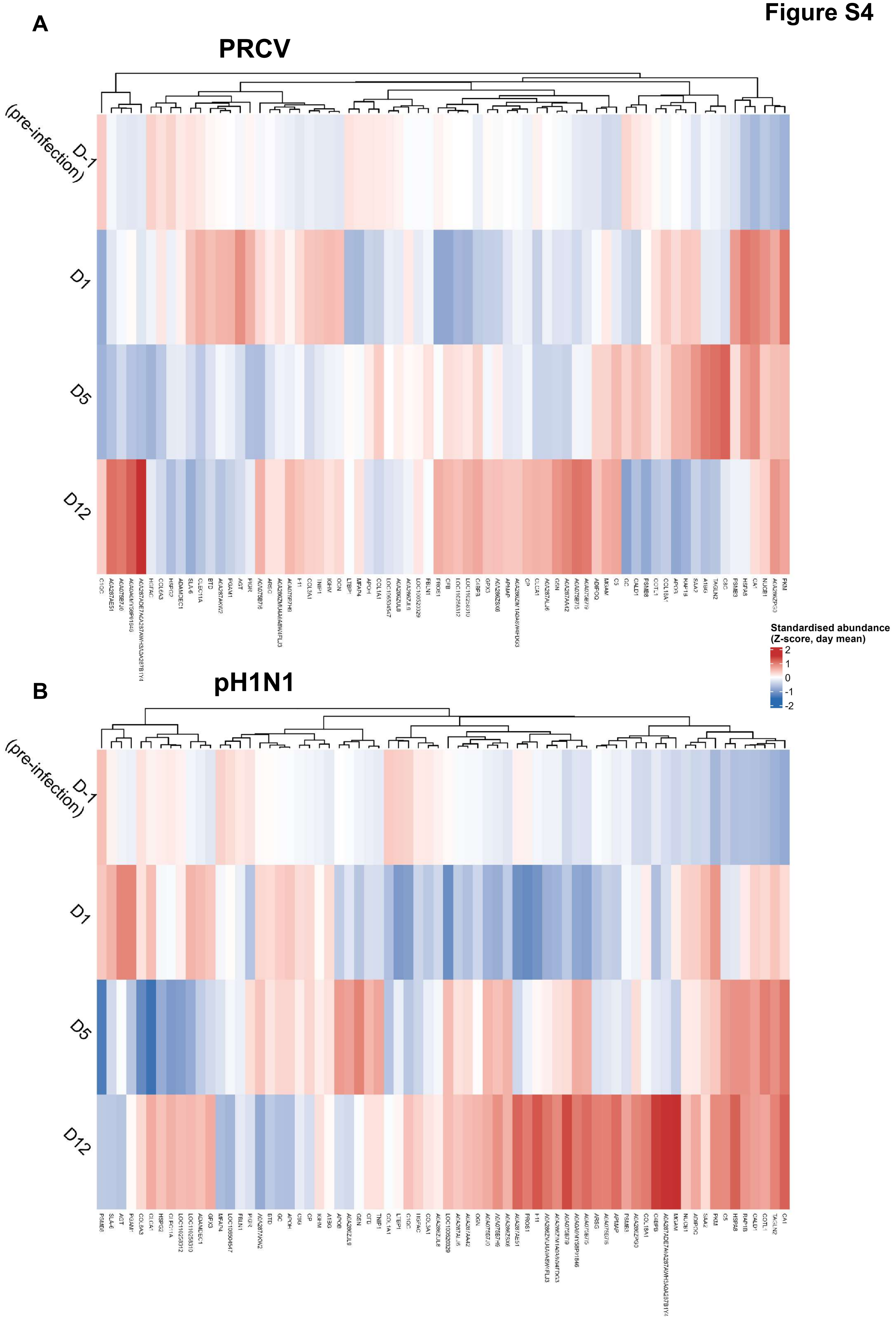
A
PRCV
D-1 (pre-infection)
D1
D5
D12
Standardised abundance
(Z-score, day mean)
2
1
0
-1
-2
B
pH1N1
D-1 (pre-infection)
D1
D5
D12

### Figure S5

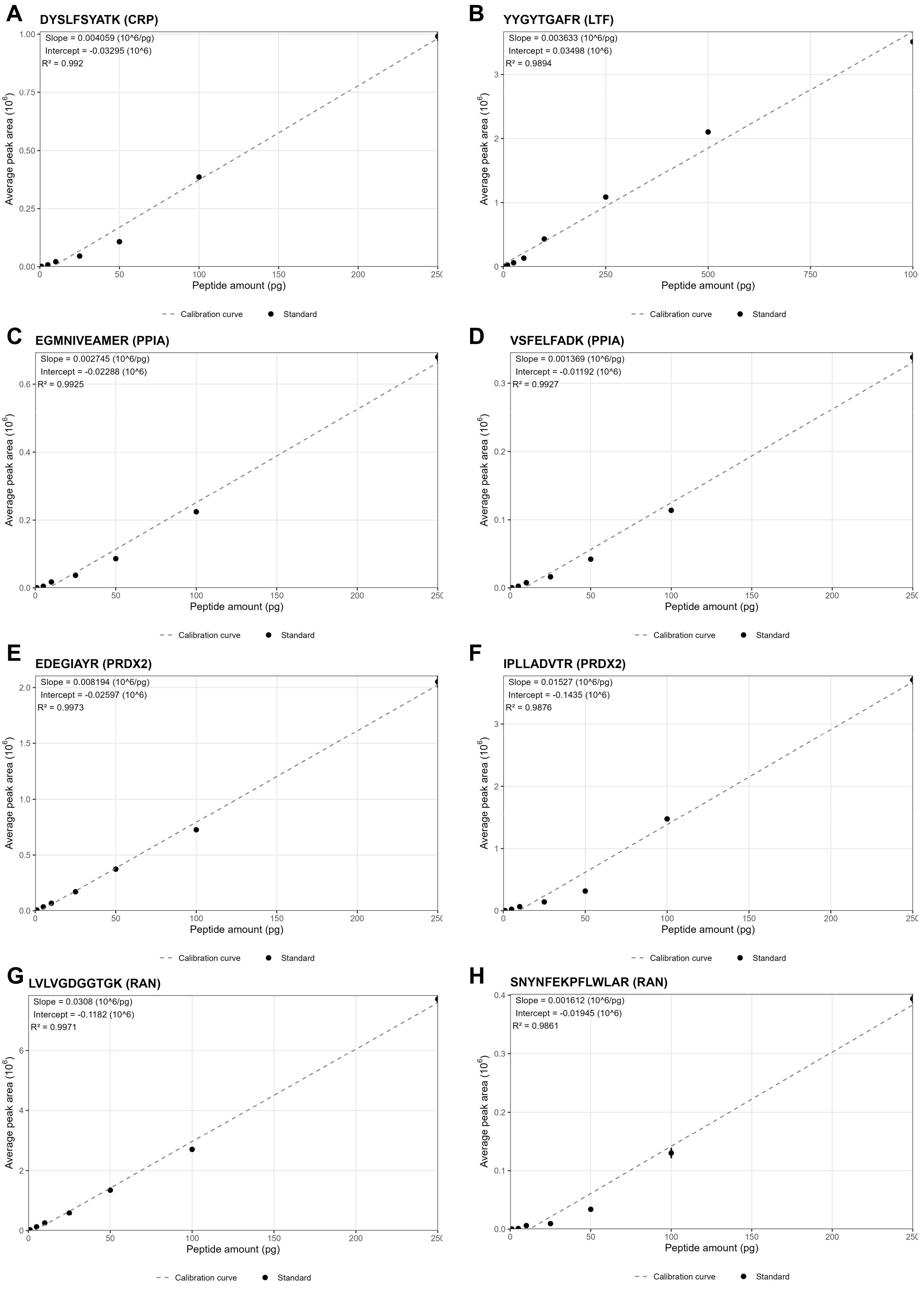
A
DYSLFSYATK (CRP)
Slope = 0.004059 (10^6/pg)
Intercept = -0.03295 (10^6)
R² = 0.992
B
YYGYTGAFR (LTF)
Slope = 0.003633 (10^6/pg)
Intercept = 0.03498 (10^6)
R² = 0.9894
C
EGMNIVEAMER (PPIA)
Slope = 0.002745 (10^6/pg)
Intercept = -0.02288 (10^6)
R² = 0.9925
D
VSFELFADK (PPIA)
Slope = 0.001369 (10^6/pg)
Intercept = -0.01192 (10^6)
R² = 0.9927
E
EDEGIAYR (PRDX2)
Slope = 0.008194 (10^6/pg)
Intercept = -0.02597 (10^6)
R² = 0.9973
F
IPLLADVTR (PRDX2)
Slope = 0.01527 (10^6/pg)
Intercept = -0.1435 (10^6)
R² = 0.9876
G
LVLVGDGGTGK (RAN)
Slope = 0.0308 (10^6/pg)
Intercept = -0.1182 (10^6)
R² = 0.9971
H
SNYNFEKPFLWLAR (RAN)
Slope = 0.001612 (10^6/pg)
Intercept = -0.01945 (10^6)
R² = 0.9861
Average peak area (10^6)
Peptide amount (pg)
Calibration curve
Standard

### Figure S5

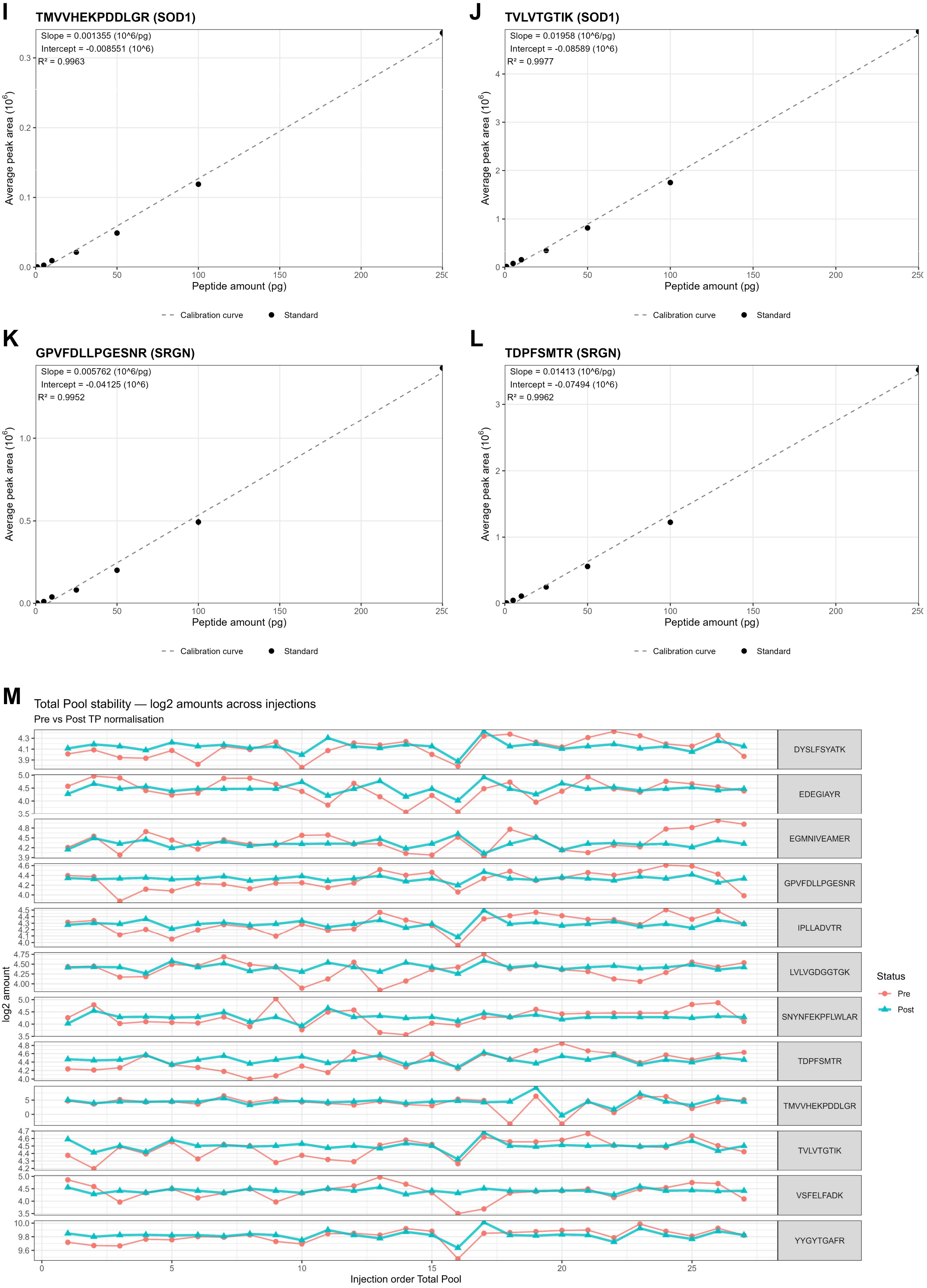

I
TMVVHEKPDDLGR (SOD1)
Slope = 0.001355 (10^6/pg)
Intercept = -0.008551 (10^6)
R² = 0.9963
J
TVLVTGTIK (SOD1)
Slope = 0.01958 (10^6/pg)
Intercept = -0.08589 (10^6)
R² = 0.9977
K
GPVFDLLPGESNR (SRGN)
Slope = 0.005762 (10^6/pg)
Intercept = -0.04125 (10^6)
R² = 0.9952
L
TDPFSMTR (SRGN)
Slope = 0.01413 (10^6/pg)
Intercept = -0.07494 (10^6)
R² = 0.9962
Average peak area ($10^6$)
Peptide amount (pg)
Calibration curve
Standard
M
Total Pool stability — log2 amounts across injections
Pre vs Post TP normalisation
DYSLFSYATK
EDEGIAYR
EGMNIVEAMER
GPVFDLLPGESNR
IPLLADVTR
LVLVGDGGTGK
SNYNFEKPFLWLAR
TDPFSMTR
TMVVHEKPDDLGR
TVLVTGTIK
VSFELFADK
YYGYTGAFR
log2 amount
Injection order Total Pool
Status
Pre
Post

**Figure S6**

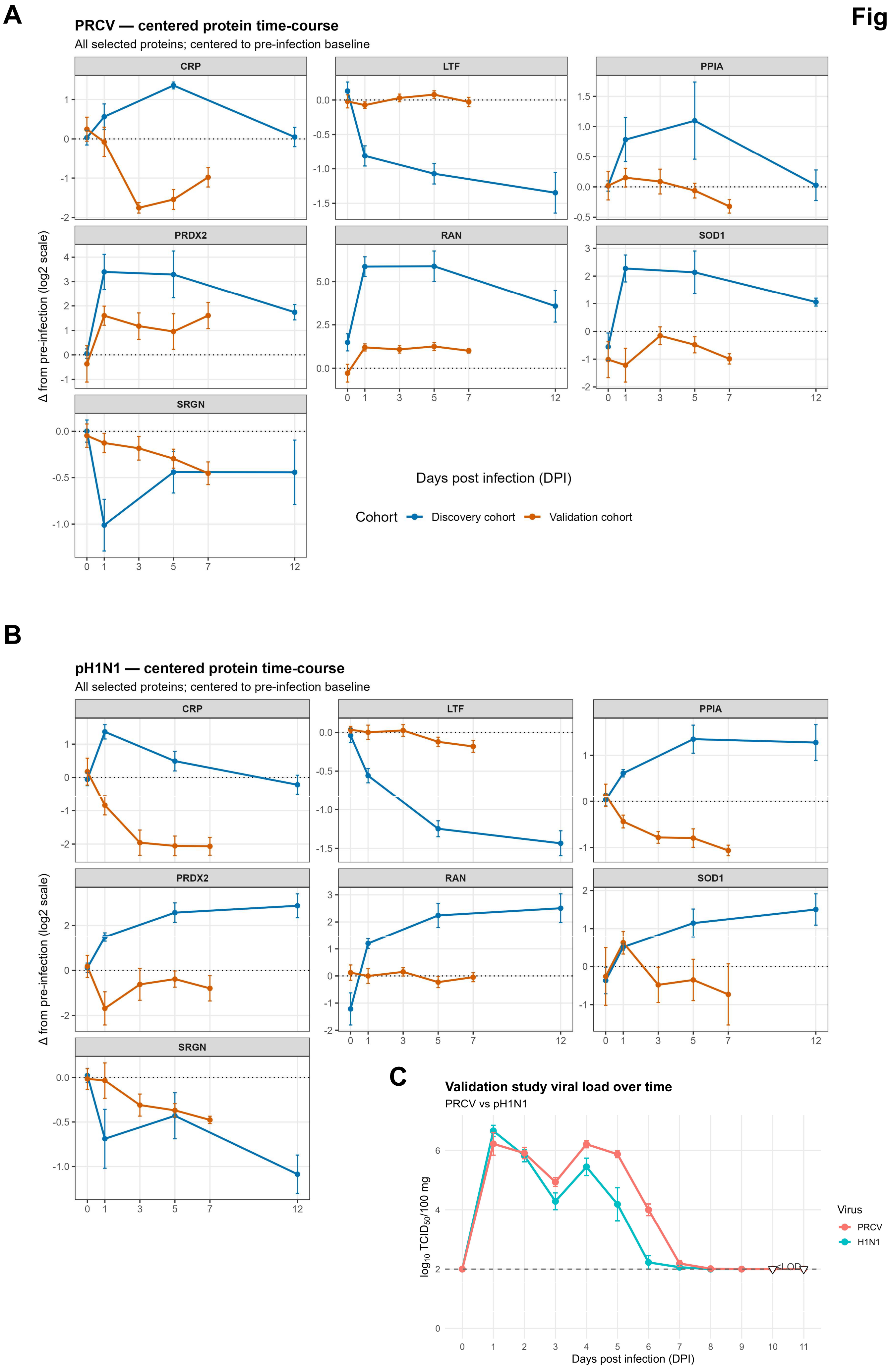

A
PRCV — centered protein time-course
All selected proteins; centered to pre-infection baseline
CRP
LTF
PPIA
PRDX2
RAN
SOD1
SRGN
Δ from pre-infection (log2 scale)
Days post infection (DPI)
Cohort
Discovery cohort
Validation cohort
B
pH1N1 — centered protein time-course
All selected proteins; centered to pre-infection baseline
CRP
LTF
PPIA
PRDX2
RAN
SOD1
SRGN
Δ from pre-infection (log2 scale)
C
Validation study viral load over time
PRCV vs pH1N1
log10 TCID50/100 mg
Days post infection (DPI)
<LOD
Virus
PRCV
H1N1

### Figure S7

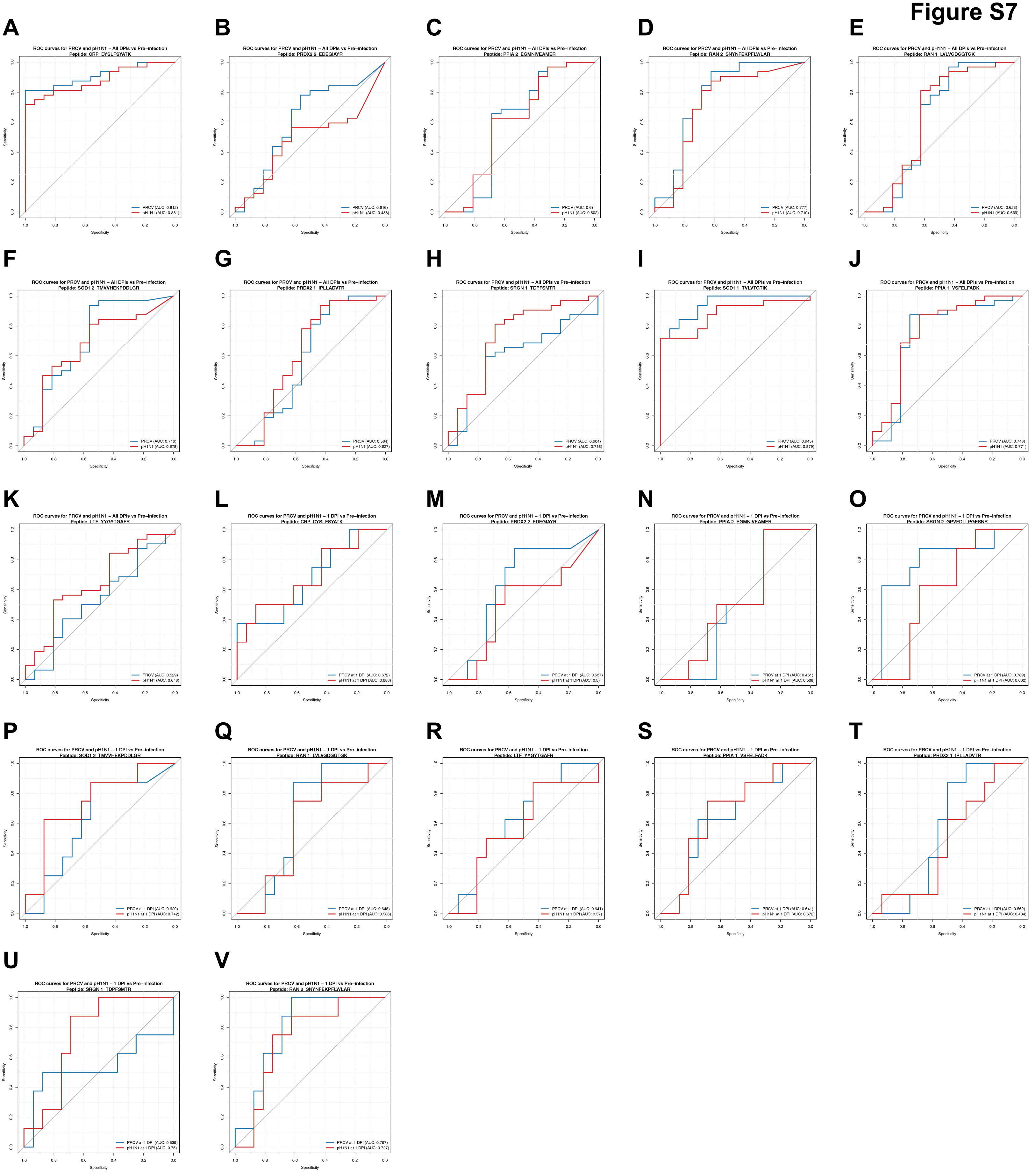
A
ROC curves for PRCV and pH1N1 – All DPIs vs Pre–infection
Peptide: CRP_DYSLFSYATK
PRCV (AUC: 0.912)
pH1N1 (AUC: 0.881)
B
ROC curves for PRCV and pH1N1 – All DPIs vs Pre–infection
Peptide: PRDX2 2_EDEGIAYR
PRCV (AUC: 0.616)
pH1N1 (AUC: 0.488)
C
ROC curves for PRCV and pH1N1 – All DPIs vs Pre–infection
Peptide: PPIA 2_EGMNIVEAMER
PRCV (AUC: 0.6)
pH1N1 (AUC: 0.602)
D
ROC curves for PRCV and pH1N1 – All DPIs vs Pre–infection
Peptide: RAN 2_SNYNFEKPFLWLAR
PRCV (AUC: 0.777)
pH1N1 (AUC: 0.719)
E
ROC curves for PRCV and pH1N1 – All DPIs vs Pre–infection
Peptide: RAN 1_LVLVGDGGTGK
PRCV (AUC: 0.623)
pH1N1 (AUC: 0.639)
F
ROC curves for PRCV and pH1N1 – All DPIs vs Pre–infection
Peptide: SOD1 2_TMVVHEKPDDLGR
PRCV (AUC: 0.716)
pH1N1 (AUC: 0.678)
G
ROC curves for PRCV and pH1N1 – All DPIs vs Pre–infection
Peptide: PRDX2 1_IPLLADVTR
PRCV (AUC: 0.584)
pH1N1 (AUC: 0.627)
H
ROC curves for PRCV and pH1N1 – All DPIs vs Pre–infection
Peptide: SRGN 1_TDPFSMTR
PRCV (AUC: 0.604)
pH1N1 (AUC: 0.738)
I
ROC curves for PRCV and pH1N1 – All DPIs vs Pre–infection
Peptide: SOD1 1_TVLVTGTIK
PRCV (AUC: 0.945)
pH1N1 (AUC: 0.879)
J
ROC curves for PRCV and pH1N1 – All DPIs vs Pre–infection
Peptide: PPIA 1_VSFELFADK
PRCV (AUC: 0.748)
pH1N1 (AUC: 0.771)
K
ROC curves for PRCV and pH1N1 – All DPIs vs Pre–infection
Peptide: LTF_YYGYTGAFR
PRCV (AUC: 0.529)
pH1N1 (AUC: 0.648)
L
ROC curves for PRCV and pH1N1 – 1 DPI vs Pre–infection
Peptide: CRP_DYSLFSYATK
PRCV at 1 DPI (AUC: 0.672)
pH1N1 at 1 DPI (AUC: 0.688)
M
ROC curves for PRCV and pH1N1 – 1 DPI vs Pre–infection
Peptide: PRDX2 2_EDEGIAYR
PRCV at 1 DPI (AUC: 0.637)
pH1N1 at 1 DPI (AUC: 0.5)
N
ROC curves for PRCV and pH1N1 – 1 DPI vs Pre–infection
Peptide: PPIA 2_EGMNIVEAMER
PRCV at 1 DPI (AUC: 0.461)
pH1N1 at 1 DPI (AUC: 0.508)
O
ROC curves for PRCV and pH1N1 – 1 DPI vs Pre–infection
Peptide: SRGN 2_GPVFDLLPGESNR
PRCV at 1 DPI (AUC: 0.789)
pH1N1 at 1 DPI (AUC: 0.602)
P
ROC curves for PRCV and pH1N1 – 1 DPI vs Pre–infection
Peptide: SOD1 2_TMVVHEKPDDLGR
PRCV at 1 DPI (AUC: 0.629)
pH1N1 at 1 DPI (AUC: 0.742)
Q
ROC curves for PRCV and pH1N1 – 1 DPI vs Pre–infection
Peptide: RAN 1_LVLVGDGGTGK
PRCV at 1 DPI (AUC: 0.648)
pH1N1 at 1 DPI (AUC: 0.586)
R
ROC curves for PRCV and pH1N1 – 1 DPI vs Pre–infection
Peptide: LTF_YYGYTGAFR
PRCV at 1 DPI (AUC: 0.641)
pH1N1 at 1 DPI (AUC: 0.57)
S
ROC curves for PRCV and pH1N1 – 1 DPI vs Pre–infection
Peptide: PPIA 1_VSFELFADK
PRCV at 1 DPI (AUC: 0.641)
pH1N1 at 1 DPI (AUC: 0.672)
T
ROC curves for PRCV and pH1N1 – 1 DPI vs Pre–infection
Peptide: PRDX2 1_IPLLADVTR
PRCV at 1 DPI (AUC: 0.562)
pH1N1 at 1 DPI (AUC: 0.484)
U
ROC curves for PRCV and pH1N1 – 1 DPI vs Pre–infection
Peptide: SRGN 1_TDPFSMTR
PRCV at 1 DPI (AUC: 0.539)
pH1N1 at 1 DPI (AUC: 0.75)
V
ROC curves for PRCV and pH1N1 – 1 DPI vs Pre–infection
Peptide: RAN 2_SNYNFEKPFLWLAR
PRCV at 1 DPI (AUC: 0.797)
pH1N1 at 1 DPI (AUC: 0.727)
Sensitivity
Specificity
